# Stoichiometric Encoding of Amplification Kinetics Enables High-Level Multiplexing in Single-Channel TaqMan Real-Time PCR

**DOI:** 10.64898/2026.11.05.764713

**Authors:** Louis Kreitmann, Kenny Malpartida-Cardenas, Ye Mao, Karolina Sobala, Zexuan Zhao, San Chun Hin, Anirudhha Hazarika, Ke Xu, Luca Miglietta, Zara Breese, Alison H. Holmes, Karen Brengel-Pesce, Laurent Drazek, Jesus Rodriguez-Manzano

**Author notes:** LK, KMC, and YM contributed equally to this work.

## Abstract

Multiplexing in TaqMan real-time PCR (qPCR) is limited to four-six targets per reaction by fluorophore spectral overlap. Amplification curve analysis (ACA) can discriminate several targets within one fluorescence channel from amplification curves (AC) kinetic features, but has been confined mainly to digital qPCR (dqPCR) and is sensitive to domain shifts between instruments and specimen types. Here, we co-design biochemical encoding of target-specific information into ACs and their computational decoding to achieve single-channel 8-plex TaqMan qPCR on a standard thermocycler. Using asymmetric PCR, we show that forward primer, reverse primer and probe concentrations act as a programmable encoding mechanism, generating a 27-condition design space of reproducible non-canonical ACs, conserved from digital to bulk qPCR. Applying this stoichiometric encoding to an 8-plex central nervous system pathogens panel increased inter-target separability, raising the mean silhouette score from 0.329 to 0.711. To decode ACs without labelled target-domain bulk qPCR data, we pretrained a transformer encoder by self-supervised learning on more than 170,000 unlabelled dqPCR curves and aligned source and target domains using conditional domain adversarial training. On contrived clinical specimens, the framework classified a canonical 7-plex respiratory virus panel with 83.6% accuracy (versus 67.7% for a supervised baseline) and the encoded 8-plex panel with 89.8% accuracy. Furthermore, stoichiometric encoding enabled resolution of double- and triple-target co-amplification events with 98.7% accuracy in controlled dqPCR experiments. This approach expands the information capacity of existing qPCR instrumentation without new hardware, fluorophores or oligonucleotides, offering a low-cost route to syndromic molecular diagnostics.

## 1. Introduction

Polymerase chain reaction (PCR)-based diagnostics enable accurate, rapid, and affordable detection of molecular targets across a wide range of clinical and analytical applications. In syndromic testing of infections, however, many different targets must be interrogated simultaneously, making multiplexing capacity a major determinant of assay throughput and clinical utility (Ramanan et al., 2017; Shin et al., 2019). In conventional TaqMan real-time PCR (qPCR), single-well differentiation of multiple targets is achieved through target-specific oligonucleotide probes labelled with spectrally distinct fluorophores (Wittwer et al., 2001). This approach is inherently constrained by spectral overlap and by the limited number of optical channels available on standard thermocyclers, typically restricting conventional assays to approximately four to six simultaneously distinguishable fluorescence signals (Kreitmann, Miglietta, et al., 2023).

In TaqMan qPCR, the fluorescence signal recorded over successive amplification cycles forms a characteristic sigmoidal amplification curve (AC). AC features reflect the kinetics and thermodynamics of the underlying amplification reaction (Heid et al., 1996) and represent an underexploited source of information that can be leveraged for target discrimination beyond conventional fluorescence readouts. Amplification curve analysis (ACA) uses kinetic signatures embedded within ACs as molecular fingerprints, enabling multiple targets to be discriminated within a single fluorescence channel using data-driven classifiers (Moniri, Miglietta, Malpartida-Cardenas, et al., 2020; Moniri, Miglietta, Holmes, et al., 2020; Miglietta, Moniri, et al., 2021). Complementary strategies have sought to improve ACA performance by increasing the intrinsic separability of ACs. Smart-Plexer is an experimental and computational framework that illustrates this principle by optimising assay selection for ACA based on inter-curve distances (Miglietta, Y. Chen, et al., 2023; Xu et al., 2025). However, Smart-Plexer selects assays according to differences between their existing canonical ACs rather than deliberately engineering new target-specific curve morphologies. Amplitude modulation provides an alternative encoding strategy by varying probe concentrations to generate target-specific differences in final fluorescence intensity (FFI), but relies predominantly on a single scalar feature rather than the full temporal structure of the AC (Rajagopal et al., 2019).

To our knowledge, no previous approach has deliberately engineered the full temporal morphology of the AC as an information-rich signal for target identification within a single fluorescence channel. Here, we hypothesised that reaction stoichiometry could provide a programmable encoding mechanism: by varying the relative concentrations of the forward primer, reverse primer and hydrolysis probe, target-specific kinetic signatures could be robustly encoded into AC morphology and subsequently decoded using data-driven methods.

Decoding such engineered signals nevertheless presents an important computational challenge. AC morphology varies substantially with template concentration, sample matrix, experimental platform, and other reaction conditions. These sources of variation generate domain shifts that can substantially impair the performance of classifiers trained on labelled source data when applied to experimentally distinct target datasets. Domain adaptation methods, including adversarial approaches such as conditional domain adversarial networks (CDAN), have been proposed to address this issue (M. Long et al., 2018), but previous applications to PCR data have primarily focused on digital qPCR (dqPCR), where large labelled datasets can be generated (Mao et al., 2023). In bulk qPCR, generating sufficient labelled data remains a major challenge. To overcome these limitations, we hypothesised that self-supervised representation learning could extract robust representations of amplification kinetics from large unlabelled datasets, while adversarial domain adaptation could subsequently align these representations across experimental domains.

Here, we establish stoichiometric encoding of AC morphology as a strategy for high-level multiplex molecular detection in single-channel TaqMan qPCR. We first map a 27-condition design space generated by varying forward primer, reverse primer and probe concentrations, and demonstrate that the resulting kinetic signatures are conserved between bulk and digital qPCR. We then apply this encoding strategy to an 8-plex assay and quantify its effect on inter-target separability relative to a conventional non-encoded panel. Third, we combine amplification-specific self-supervised representation learning with conditional adversarial domain adaptation to decode the engineered signals in bulk qPCR. Finally, we investigate whether the information contained within these engineered ACs can be used to distinguish controlled double-, and triple-target co-amplification events.

Together, this co-design of molecular signal encoding and computational decoding provides a framework for increasing the multiplexing capacity of conventional real-time PCR without requiring additional fluorescence channels or modification of existing instrumentation. It offers a practical and scalable strategy to complement established multiplex PCR solutions, increasing diagnostic throughput while reducing platform complexity, footprint, and cost.

## 2. Material & Methods

### 2.1 Reagents, oligonucleotides and instruments

All primers and double-quenched TaqMan probes were synthesised by Integrated DNA Technologies (IDT, United States), and reactions used 2*×* IDT PrimeTime Gene Expression Master Mix (catalogue number 1055772). Bulk qPCR reactions had a final volume of 10 *µ*L and contained 5 *µ*L 2*×* master mix, 1 *µ*L 20*×* GE, 1 *µ*L 10*×* primer mix, 1 *µ*L DNA template (or NTC), and nuclease-free water. Reactions were performed on a QuantStudio 5 Real-Time PCR System (Thermo Fisher Scientific, United States), with an initial hot-start step at 95 ^*°*^C for 3 min followed by 50–60 cycles of 95 ^*°*^C for 15 s and 60 ^*°*^C for 30–45 s. Real-time digital PCR (dqPCR) was performed on a Biomark HD system (Fluidigm Corporation, South San Francisco, CA, USA) using qdPCR 37K integrated fluidic circuits. dqPCR reactions had a final volume of 6 *µ*L and contained 3 *µ*L 2*×* master mix, 0.6 *µ*L 20*×* GE, 0.6 *µ*L 10*×* primer mix, 1.8 *µ*L DNA template (or NTC), and nuclease-free water; 4.5 *µ*L of reaction mix was loaded into each IFC inlet. Cycling conditions were identical to those used for bulk qPCR. Template, primer and probe sequences and full experimental details are provided in the Supplementary Material.

### 2.2 7-plex TaqMan panel for respiratory viruses

We used a previously published 7-plex TaqMan panel targeting the hexon gene of human adenovirus (HAdV) and the nucleocapsid (N) gene of human coronaviruses OC43 (HCoV-OC43), HKU1 (HCoV-HKU1), 229E (HCoV-229E), NL63 (HCoV-NL63), Middle East respiratory syndrome-related coronavirus (MERS-CoV) and severe acute respiratory syndrome coronavirus 2 (SARS-CoV-2). This combination was selected by the Smart-Plexer algorithm as the combination with the highest separability index—the greatest distance between ACs—among 4,608 possible 7-plex combinations drawn from 24 singleplex assays (Miglietta, Y. Chen, et al., 2023).

### 2.3 8-plex panel for central nervous system infections

The 8-plex central nervous system (CNS) pathogen panel includes eight targets: the *ply* gene of *Streptococcus pneumoniae* (Spne), the *ctrA* gene of *Neisseria meningitidis* (Nmen), the *hly* gene of *Listeria monocytogenes* (Lmon), the gD gene of herpes simplex virus 1 (HSV1), the gG gene of herpes simplex virus 2 (HSV2), the *bexA* gene of *Haemophilus influenzae* (Hinf), the *cfb* gene of *Streptococcus agalactiae* (Saga), and the 18S rDNA target of *Cryptococcus neoformans* (Cneo). TaqMan assays were selected from previously published data (Corless et al., 2001; Tzanakaki et al., 2005; Messmer et al., 1995; Albuquerque et al., 2019; Wang et al., 2020; Zhou et al., 2016; Bøving et al., 2009; F. Long et al., 2019; Khumalo et al., 2017; Shi et al., 2016; Markoulatos et al., 2001; Lusia Leal et al., 2008) and validated in singleplex under symmetric conditions using serial dilutions of synthetic DNA to assess analytical sensitivity and AC quality. Assays were then combined incrementally into a non-encoded 8-plex panel, and AC shapes were confirmed to be unchanged between the singleplex and multiplex settings.

We then introduced stoichiometric encoding by assigning each target a condition drawn from the 3 *×* 3 *×* 3 array of asymmetric PCR designs described below. Serial dilutions of synthetic DNA were run in multiplex and AC morphologies were assessed; assay conditions were amended where the observed ACs did not match the intended encoding, and iterative optimisation was continued until the final encoded 8-plex CNS panel was obtained (Table S3, Figure S2a). dqPCR experiments confirmed that the encoded morphologies were conserved in partitioned reactions (Figure S2b). The separability of ACs in the non-encoded and encoded panels was compared using two-dimensional principal component analysis (2D-PCA; Figures 1g and 1h).

**Figure 1.**
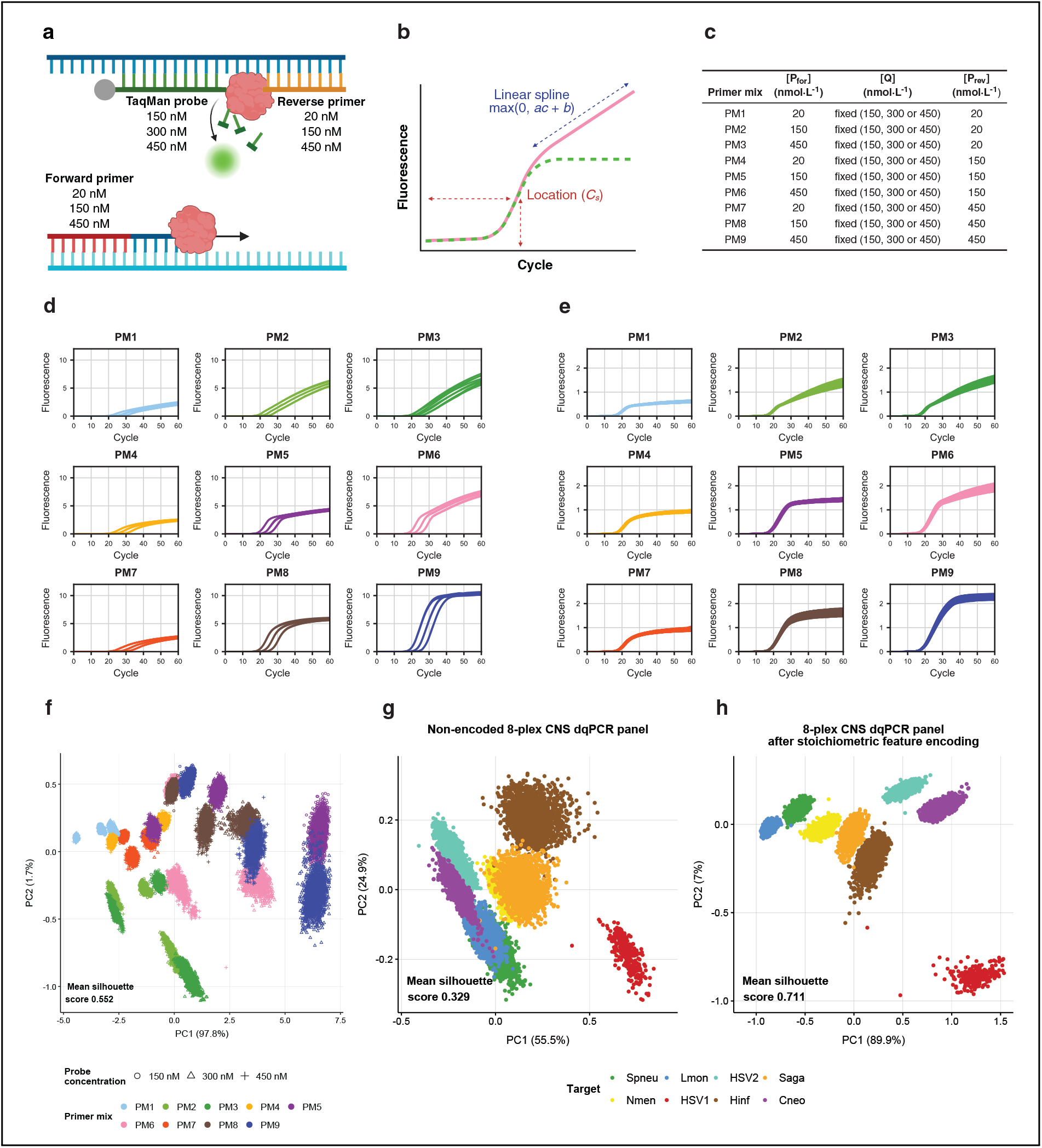
Stoichiometric encoding enables programmable modulation of AC morphology and increases target separability. **a. Experimental design of asymmetric TaqMan PCR**. The concentrations of the forward (*P*_for_) and reverse (*P*_rev_) primers are varied independently across three levels (20, 150 and 450 nmol*·*L^*−*1^), while the probe *Q* is tested at three concentrations (150, 300, and 450 nmol*·*L^*−*1^). This generates a three-by-three-by-three design array yielding distinct AC morphologies. **b. Generation of non-canonical ACs**. Amplification proceeds symmetrically until the limiting primer is depleted (dashed green line), after which single-stranded amplification from the excess primer drives a linear increase in fluorescence (solid pink line). This transition results in non-canonical AC shapes that can be exploited for target discrimination in multiplex assays. **c. Composition of primer mixes**. Nine primer mixes are generated by combining *P*_for_ and *P*_rev_ at three concentrations (20, 150, and 450 nmol*·*L^*−*1^) in all pairwise combinations. Within each set of nine PMs, the probe concentration is fixed at 150, 300, or 450 nmol*·*L^*−*1^, leading to a three-by-three-by-three array of modulation strategies. **d. Asymmetric bulk qPCR experiment with three concentrations of the HCoV-229E template**. Bulk qPCR experiments were conducted with three concentrations (10^6^, 10^5^, and 10^4^ copies + NTCs) of the HCoV-229E template in triplicate. Distinct AC shapes are observed across the three-by-three matrix of primer ratios at fixed [*Q*] = 450 nmol *·* L^*−*1^. Diagonal conditions (PM1, PM5, PM9) correspond to symmetric assays with equal primer concentrations, whereas off-diagonal conditions produce non-canonical curves characterised by an initial exponential phase followed by linear fluorescence. The transition point between these phases depends on the ratio of limiting to excess primer. The matrix is asymmetric, reflecting strand-specific probe binding in TaqMan PCR. **e. dqPCR experiment with the HCoV-229E template**. ACs obtained in dqPCR (36,960 partitions) show patterns consistent with the corresponding bulk qPCR data, confirming that stoichiometric feature encoding is preserved in partitioned reactions. **f. 2D-PCA visualisation of the feature design space**. 2D-PCA plot of dqPCR ACs obtained with the HCoV-229E template (62,370 ACs in total obtained in three separate experiments) across 27 distinct designs, coloured by primer mix with marker style indicating probe concentration, reveals clear clustering according to PM conditions. **g, h. 2D-PCA of dqPCR ACs obtained with the 8-plex CNS infection panel**. Panel g shows the non-encoded baseline assay (16,554 ACs), whereas panel h shows the stoichiometrically encoded assay (16,460 ACs). Both datasets were acquired in the same run on a single 37K integrated fluidic circuit (Fluidigm Corporation, South San Francisco, CA, USA). Stoichiometric encoding markedly increased class separability, with mean silhouette score increasing from 0.329 to 0.711.

### 2.4 Stoichiometric design space

Nine primer mixes (PM1–PM9) were generated by combining the forward (*P*_for_) and reverse (*P*_rev_) primers at 20, 150 or 450 nmol*·*L^*−*1^ in all nine pairwise combinations (Figure 1c). Within each set of nine primer mixes, the probe (*Q*) concentration was fixed at 150, 300 or 450 nmol*·*L^*−*1^, giving a 3 *×* 3 *×* 3 array of 27 conditions. The diagonal conditions PM1, PM5 and PM9 correspond to symmetric PCR, with equal forward and reverse primer concentrations. Because the hydrolysis probe binds only one strand of the amplicon, single-stranded product generated after depletion of the limiting primer is detected only when the excess primer extends the strand complementary to the probe; the primer matrix is therefore not symmetric with respect to the exchange of *P*_for_ and *P*_rev_.

### 2.5 Contrived specimens and ethics

Because residual clinical specimens were unavailable for several targets, we generated contrived specimens by spiking synthetic DNA templates into human DNA extracted from ten residual anonymised respiratory swabs using the QIAsymphony^®^ DSP Virus/Pathogen Midi Kit (Cat. No. 937055, QIA-GEN). The swabs had previously tested negative for an extended panel of respiratory viruses and were obtained through the virology laboratory at Imperial College Healthcare NHS Trust under IRAS Project ID 289096. DNA extracts from the ten swabs were pooled and used at an approximate human DNA concentration of 4 *×* 10^4^ genome copies per *µ*L. The same human-DNA background was used for the respiratory and CNS panels to provide a consistent biological matrix across experiments and isolate assay- and platform-related effects. Contrived-specimen datasets were treated as unlabelled during model training to reflect a deployment setting in which target-domain labels are unavailable, whereas their labels were used only for evaluation of classification performance; synthetic DNA datasets were used with known labels.

### 2.6 Template concentrations

dqPCR was performed at approximately 10^6^, 10^5^ and 10^4^ copies per panel inlet, corresponding to expected mean occupancies of *λ* = 142, 14.2 and 1.4 copies per partition, respectively, with no-template controls (NTCs). Bulk qPCR was performed across seven concentrations (10^7^ to 1 copy per reaction), together with NTCs; SARS-CoV-2 was limited to 10^5^ copies per reaction for biosecurity reasons.

### 2.7 Pre-processing of amplification data

ACs from both dqPCR and bulk qPCR were pre-processed before being used as input to machine-learning models. Curves showing no amplification, including NTCs and negative dqPCR partitions, were discarded. Baselines were set to zero using an in-house cycle threshold (*C*_*t*_)-based correction procedure. For each AC, the *C*_*t*_ was identified from the maximum second derivative of the fluorescence signal, and the mean background fluorescence over a window of 5–10 cycles preceding the *C*_*t*_ was subtracted from the entire curve; resulting negative fluorescence values were floored to zero. To maintain comparable fluorescence scales across experimental runs and platforms, each experiment included a synthetic DNA positive control, and fluorescence values were normalised to its mean final fluorescence intensity. Finally, curves were smoothed using a Savitzky–Golay filter with a window length of 15 cycles and polynomial order 3 (Luo et al., 2005). Further details are provided in the Supplementary Material.

### 2.8 Self-supervised pretraining with BYOL

Bootstrap Your Own Latent (BYOL) is a self-supervised representation-learning framework that does not require explicit negative pairs (Grill et al., 2020). It comprises an online network, with parameters *θ* and consisting of an encoder *f*_*θ*_, projector *g*_*θ*_ and predictor *q*_*θ*_, together with a target network with parameters *ξ* (Figure 2b). Two independently augmented views of the same input are passed through the online and target networks. The online network is trained to predict the target-network projection of the alternative view by minimising the squared distance between *l*_2_-normalised predictions and target projections, symmetrised over the two views. The target-network parameters are updated as an exponential moving average of the online-network parameters.

**Figure 2.**
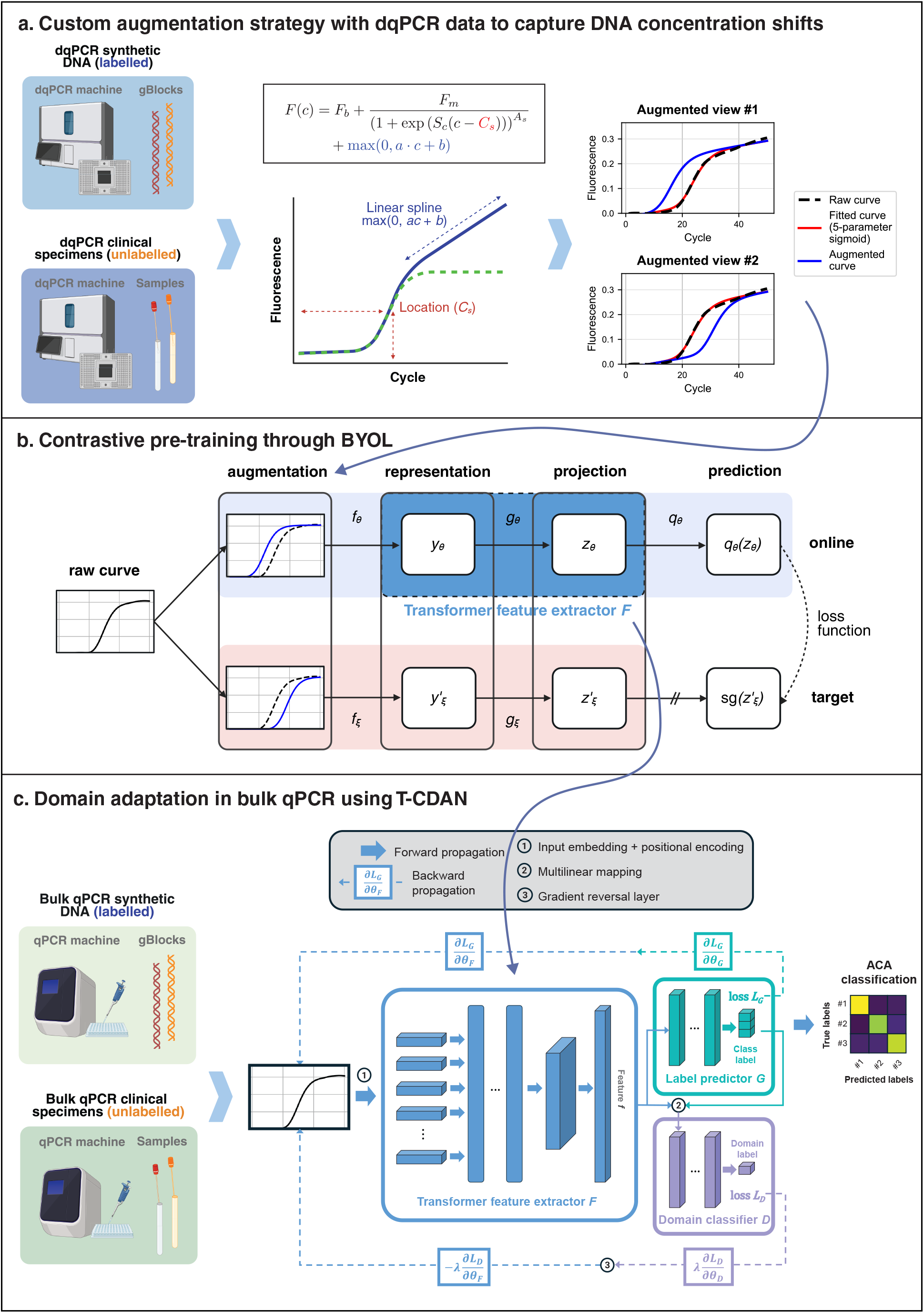
Self-supervised representation learning and adversarial domain adaptation for ACA. **a. Custom augmentation procedure using non-canonical ACs in dqPCR to capture concentration shifts of input DNA**. Each raw AC from the dqPCR contrived clinical dataset is fitted using a composite model combining a 5-parameter sigmoid and hinge linear component to capture asym-metric amplification dynamics (red). Augmented views (blue) are then generated by selectively varying the location parameter (*C*_*s*_) while keeping other parameters fixed (see Methods for details on the augmentation algorithm). This procedure simulates realistic shifts in amplification timing associated with differences in input DNA concentration, while preserving the intrinsic target-specific kinetic and thermodynamic characteristics of the curve. Two independently augmented views of the same curve are used as positive pairs for self-supervised learning. **b. Self-supervised pretraining using BYOL**. The two augmented views are processed independently through an online network (encoder *f*_*θ*_, projector *g*_*θ*_, predictor *q*_*θ*_) and a target network (encoder *f*_*ξ*_, projector *g*_*ξ*_). In this work, *f*_*θ*_ is implemented as a stack of transformer encoder blocks. The online network is trained to predict the target network projection by minimising a similarity loss between normalised projections, while the target network parameters are updated as an exponential moving average of the online network weights. After training, the encoder *f*_*θ*_ provides a robust representation of ACs. Adapted from Grill et al., 2020. **c. Domain adaptation using T-CDAN**. The T-CDAN framework comprises three main components: a feature extractor *F*, a label predictor *G*, and a domain classifier *D*. The feature extractor *F* maps a standardised AC **x** to a feature representation ***f***. In the final classification pipeline, the encoder *f*_*θ*_ pretrained through BYOL is used to initialise *F*, enabling the model to leverage previously learnt representations of ACs. The label predictor is a standard feed-forward neural network trained to classify source-domain data into target labels, while *D* is a two-layer perceptron trained to distinguish between source and target domains. During training, the label prediction loss *L*_*G*_ and the domain classification loss *L*_*D*_ are optimised adversarially, enabling *F* to learn domain-invariant features of ACs. Numbers 1–3 indicate forward and backward propagation paths. Reproduced from Mao et al., 2023, with permission from IEEE©.

Whereas the original BYOL architecture was developed for images, here the inputs were ACs represented as one-dimensional sequences of length *c*, corresponding to the number of PCR cycles, and the encoder was a one-dimensional transformer. The augmented views *v* = *t*(*x*) and *v*^*′*^ = *t*^*′*^(*x*) were generated using the AC-specific augmentation procedures described below. Following pretraining, the online encoder *f*_*θ*_ was used to initialise the T-CDAN feature extractor *F*, which was subsequently updated during domain-adaptation training. Detailed loss definitions, architecture and training hyperparameters are provided in the Supplementary Material.

### 2.9 Augmentation strategies

Because standard image augmentations are not applicable to ACs, we developed a domain-specific augmentation strategy based on parametric models of amplification kinetics (Swillens et al., 2008; Miglietta, Xu, et al., 2022). Canonical ACs were modelled using a five-parameter sigmoid:

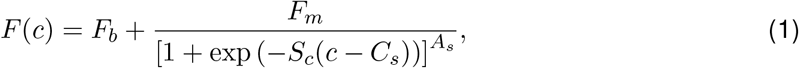

where *c* is the PCR cycle number, *F* (*c*) is the fluorescence at cycle *c, F*_*b*_ is the baseline fluorescence, *F*_*m*_ is the maximal fluorescence, *C*_*s*_ is a location parameter corresponding to the point of inflexion, *S*_*c*_ *>* 0 controls the slope of the exponential portion of the curve, and *A*_*s*_ controls asymmetry of the sigmoid (Spiess et al., 2008). Each AC was fitted by nonlinear least squares (Moré, 1978; Branch et al., 1999) using `scipy.optimize.curve fit` (Virtanen et al., 2020), yielding the parameter set *S* = *{F*_*b*_, *F*_*m*_, *C*_*s*_, *S*_*c*_, *A*_*s*_*}*.

Augmented views were generated by altering only the location parameter *C*_*s*_, while retaining all other fitted parameters. For each augmented view,

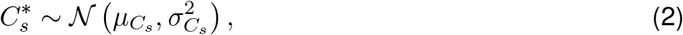

where 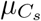 and 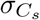 are the empirical mean and standard deviation of fitted *C*_*s*_ values in the unlabelled dqPCR dataset. The sigmoid was then re-evaluated using 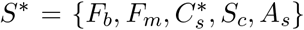. Two independently generated views of each raw AC formed a positive pair for BYOL pretraining. This procedure introduces variation in amplification timing associated with differences in input DNA concentration while preserving the remaining kinetic characteristics of the curve (Rutledge and Côté, 2003; Gevertz et al., 2005).

For the stoichiometrically encoded 8-plex CNS panel, the canonical sigmoid model was extended because depletion of the limiting primer during asymmetric PCR produces a secondary approximately linear increase in fluorescence. These ACs were therefore fitted using a sigmoid-plus-linear model:

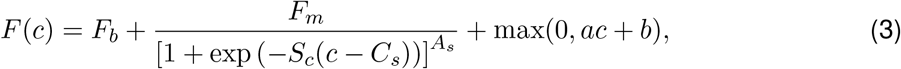

where *F*_*b*_, *F*_*m*_, *C*_*s*_, *S*_*c*_ and *A*_*s*_ are defined as above, and max(0, *ac* + *b*) is a hinge-linear component with slope *a* and intercept *b* capturing late-cycle single-stranded amplification. Augmented views were generated as above by resampling *C*_*s*_ while keeping all other parameters fixed.

### 2.10 Domain adaptation using T-CDAN

T-CDAN comprises three components: a transformer-based feature extractor *F*, a label predictor *G* and a conditional domain classifier *D* (Figure 2c) (Mao et al., 2023). During training, labelled source-domain data *S* (synthetic DNA) and unlabelled target-domain data *T* (contrived specimens) are passed through *F*. Source-domain features are used by *G* to predict target identity, whereas features from both domains are supplied to *D* to predict their domain of origin. Adversarial training encourages *F* to generate representations that retain information relevant to target classification while reducing the distributional difference between source and target domains (M. Long et al., 2018; Mirza and Osindero, 2014; Yu et al., 2021).

ACs are supplied to *F* as *c*-dimensional vectors. Each standardised curve is mapped to an *N*_*d*_-dimensional latent representation by a learnable embedding layer, combined with positional encoding, and processed by a stack of transformer blocks comprising multi-head self-attention and position-wise multilayer perceptrons. The resulting representation **f** is supplied to the label predictor *G*, which produces class logits **g** and is optimised using categorical cross-entropy loss *ℒ* _*G*_. The conditional domain classifier receives the multilinear representation

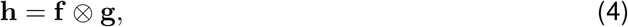

and is trained using the domain-classification loss *ℒ* _*D*_. The overall adversarial objective is

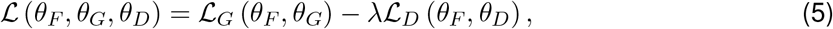

where *θ*_*F*_, *θ*_*G*_ and *θ*_*D*_ denote the parameters of *F, G* and *D*, respectively. The contribution of the domain-adversarial objective, controlled by *λ*, was progressively annealed during training. This adversarial optimisation encourages features extracted by *F* to maximise the ability of *G* to discriminate target classes while minimising the ability of *D* to discriminate experimental domains. Detailed network architecture, loss definitions and training hyperparameters are provided in the Supplementary Material.

### 2.11 Co-amplification experiments

To investigate multi-target amplification, three targets from the encoded CNS panel*—Streptococcus pneumoniae* (Spne), *Listeria monocytogenes* (Lmon), and *Streptococcus agalactiae* (Saga)—were tested individually, in all three pairwise combinations, and in the triple-target combination. Oligonu-cleotide concentrations were identical to those used in the encoded 8-plex panel (Table S4). Synthetic DNA templates were used at 10^6^, 10^5^ and 10^4^ copies per reaction, with equal input concentrations for each target in mixed reactions; bulk qPCR and dqPCR were otherwise performed as described in the Supplementary Material. The separability of the seven amplification conditions was visualised by PCA. Classification was evaluated using a *k*-nearest neighbours (kNN) classifier (*k* = 5) with grouped cross-validation, using IFC inlets as groups. kNN was used rather than the transformer-based pipeline because all seven classes were acquired within a single dqPCR domain and no domain shift had to be corrected (Moniri, Miglietta, Malpartida-Cardenas, et al., 2020; Moniri, Miglietta, Holmes, et al., 2020).

### 2.12 Evaluation protocol and statistics

Domain-adaptation experiments used a transductive setting: the unlabelled target-domain ACs used for adversarial alignment were drawn from the same target dataset subsequently used for evaluation, but their target labels were withheld throughout adaptation and used only for performance assessment. Each deep-learning model was trained once, and classification performance is reported as the observed accuracy on the corresponding evaluation dataset. No formal statistical comparisons between models were performed.

No statistical methods were used to predetermine sample size. Experiments were not randomised and investigators were not blinded. Wet-lab experiments were performed with at least two independent replicates unless otherwise stated. Analyses were performed in Python using SciPy (Virtanen et al., 2020), scikit-learn (Pedregosa et al., 2011) and PyTorch (Paszke et al., 2019).

### 2.13 Feature visualisation

We used two-dimensional principal-component analysis (2D-PCA) to visualise the structure of AC data and assess inter-class separability. PCA was applied to standardised, post-processed ACs, and the first two principal components were used for visualisation, with each point representing one AC and coloured according to target class. Cluster separability was quantified using the mean silhouette score calculated in the two-dimensional PCA space (Rousseeuw, 1987).

We additionally used two-dimensional t-distributed stochastic neighbour embedding (t-SNE) to visualise learnt feature representations across model configurations (Maaten and Hinton, 2008). Source- and target-domain feature vectors were combined and, where necessary, randomly subsampled to a maximum of 5,000 observations. Feature dimensionality was first reduced to 50 components using PCA, followed by t-SNE with a perplexity of 50, 1,000 iterations, PCA initialisation and an automatically determined learning rate; the t-SNE random seed was set to 42. Embeddings were visualised by experimental domain to assess domain alignment and by target class to assess class separability.

## 3. Results

### 3.1 Stoichiometric feature encoding enables programmable modulation of AC morphology

We first investigated whether non-canonical AC morphology could be deliberately engineered by modifying reaction stoichiometry. To this end, we used asymmetric PCR, a variation of standard PCR that preferentially amplifies one strand of a double-stranded DNA template through unequal primer concentrations (Sanchez et al., 2004) (Figure 1a). The limiting primer is supplied at a lower concentration than the excess primer; once depleted, amplification proceeds predominantly from the excess primer, generating single-stranded products that are detected by cleavage of the complementary hydrolysis probe, thereby producing non-canonical ACs (Figure 1b).

We used DYNAMIC, a software implementation of a kinetic model of single-plex TaqMan PCR acting as a digital twin (Kreitmann, Mao, et al., 2026), to predict the morphological changes produced by varying primer stoichiometry. To test these *in silico* predictions experimentally, we amplified the HCoV-229E target from our previously developed respiratory virus panel (Section 2.2) using the nine primer mixes PM1–PM9, in which *P*_for_ and *P*_rev_ were combined at three concentrations (20, 150 or 450 nmol *·* L^*−*1^) in nine pairwise combinations, with the probe fixed at 450 nmol *·* L^*−*1^ (Figure 1c).

As shown in Figure 1d, altering primer stoichiometry generated distinct AC morphologies across the 3 *×* 3 primer matrix, including non-canonical curves characterised by initial exponential amplification followed by a linear increase in fluorescence at later cycles. The transition point between the two phases depended on the ratio of limiting to excess primer, and the matrix was not symmetric with respect to *P*_for_ and *P*_rev_, reflecting strand-specific probe binding (Section 2.4). We next examined whether probe concentration provided an additional degree of control over AC morphology. Lowering [*Q*] to 300 and 150 nM produced additional distinct kinetic profiles (Figures S1a and S1b), thereby defining a 27-condition stoichiometric design space based on independent modulation of forward primer, reverse primer and probe concentrations.

We next examined whether these engineered kinetic signatures were conserved across PCR platforms. The morphological effects observed in bulk qPCR were recapitulated in partitioned reactions in dqPCR (Figures 1e, S1c and S1d). Furthermore, 2D-PCA of 62,370 ACs obtained across the 3 *×* 3 *×* 3 design space revealed well-separated and structured clusters corresponding to the different stoichiometric conditions, with a mean silhouette score of 0.552 (Figure 1f). Together, these experiments establish reaction stoichiometry as a programmable mechanism for encoding information into AC morphology in TaqMan qPCR.

### 3.2 Stoichiometric feature encoding increases signal separability in an 8-plex TaqMan assay

We next asked whether this encoding strategy could be used to increase the intrinsic separability of targets within a high-order multiplex assay. To this end, we designed an 8-plex single-channel TaqMan panel targeting eight CNS pathogens (Spne, Nmen, Lmon, Hinf, Saga, Cneo, HSV-1 and HSV-2; Section 2.3). Following validation of the individual assays under conventional symmetric conditions, targets were progressively combined into a non-encoded reference multiplex panel. We then generated an encoded panel by assigning each target a specific stoichiometric condition selected from the 27-condition design space described above. Iterative optimisation was performed to ensure that the resulting ACs reproducibly exhibited distinct target-specific morphologies (Section 2.3, Table S4).

The final encoded panel generated eight distinct and reproducible AC signatures in bulk qPCR (Figure S2a), which were again conserved in dqPCR (Figure S2b). Importantly, stoichiometric encoding substantially increased the intrinsic separation between target classes compared with the conventional non-encoded panel: in 2D-PCA of dqPCR data acquired in a single run, the mean silhouette score increased from 0.329 for the non-encoded panel (Figure 1g) to 0.711 for the encoded panel (Figure 1h). These results demonstrate that stoichiometric encoding does not merely alter AC morphology, but can be used prospectively to engineer a multiplex assay in which target-specific signals occupy more distinct regions of feature space.

### 3.3 Self-supervised representation learning and domain adaptation enable robust AC classification across experimental domains

Having established that AC morphology could be deliberately engineered to increase target separability, we next developed the computational framework capable of decoding target identity under the domain shifts encountered in bulk qPCR. Our overarching aim was to classify ACs obtained from bulk qPCR of specimens in a human-DNA background despite limited availability of labelled data obtained in that setting. The resulting framework combines self-supervised representation learning with adversarial domain adaptation (Figure 2).

We first developed and benchmarked this framework using the previously published 7-plex TaqMan respiratory virus panel (Section 2.2) (Miglietta, Y. Chen, et al., 2023). Four datasets were generated: (1) dqPCR synthetic DNA (33,345 ACs), (2) dqPCR contrived clinical specimens (59,237 ACs), (3) bulk qPCR synthetic DNA (402 ACs), and (4) bulk qPCR contrived clinical specimens (446 ACs). Together, these datasets captured both platform-related domain shift (dqPCR versus bulk qPCR) and specimen-related domain shift (synthetic DNA versus contrived clinical specimens). ACs from the four datasets are shown in Figure S3. Contrived clinical specimen datasets were treated as unlabelled, mirroring a deployment setting in which target-domain labels are unavailable, whereas labels from the synthetic DNA datasets were used for supervised training.

Self-supervised pretraining was performed using BYOL on dqPCR data treated as unlabelled (Section 2.8) (Grill et al., 2020). For each raw dqPCR AC, we generated two informative augmented “views” by fitting a five-parameter sigmoid and resampling only its location parameter (*C*_*s*_), which shifts the curve along the cycle axis and thereby emulates realistic kinetic variability associated with different input DNA concentrations while preserving target-specific thermodynamic and kinetic features (Section 2.9, Figure S4). The 5-parameter sigmoid model of amplification kinetics was appropriate for this dataset because the 7-plex TaqMan respiratory virus panel does not involve stoichiometric encoding and only comprises ‘canonical’ AC morphologies. The generated paired views were used for BYOL pretraining, in which a transformer-based online network learned representations of AC morphology by matching embeddings of the two augmented views without requiring target labels (Figure 2b). The encoder and projector components of the pretrained network were subsequently incorporated into a T-CDAN architecture, enabling unsupervised alignment of the labelled source (synthetic DNA) and unlabelled target (contrived clinical specimens) feature distributions (Section 2.10, Figure 2c).

To assess the added value of self-supervised pretraining beyond domain adaptation, we first considered classification of dqPCR contrived specimens (Figure S5a). The baseline 1D-transformer trained on dqPCR synthetic DNA (model 1.1) achieved an accuracy of 84.8% across seven targets (chance level 14.3%), demonstrating a measurable specimen-related domain shift between synthetic DNA and contrived specimens. T-CDAN alone (model 1.2) did not change performance (84.4%), suggesting that adversarial alignment alone provided limited benefit in this setting. BYOL pretraining followed by T-CDAN (model 1.3) increased accuracy to 86.8% (Figure S6), supporting the ability of self-supervised learning to extract informative representations from unlabelled ACs for subsequent target classification.

We next evaluated the framework on bulk qPCR contrived specimens, where both specimen- and platform-related domain shifts were present (Figure S5b). The dqPCR-trained baseline (model 2.1) achieved an accuracy of 51.3%, consistent with the substantial shift between PCR platforms and specimen types (Figure S7). Training directly on bulk qPCR synthetic DNA (model 2.2) increased accuracy to 67.7%, while domain adaptation (model 2.3) produced a modest further increase to 69.3%. The largest improvement was observed when self-supervised pretraining was combined with domain adaptation (model 2.4), yielding an accuracy of 83.6% across the seven targets. Together, these experiments establish self-supervised representation learning followed by adversarial domain adaptation as an effective strategy for decoding ACs in bulk qPCR when labelled target-domain data are scarce.

### 3.4 Computational decoding of biologically encoded signals in the 8-plex assay

We next tested whether the computational framework developed above could decode the deliberately engineered signals generated by the encoded 8-plex CNS panel. Four datasets were generated: (1) dqPCR synthetic DNA (81,007 ACs), (2) dqPCR contrived clinical specimens (89,507 ACs), (3) bulk qPCR synthetic DNA (489 ACs), and (4) bulk qPCR contrived clinical specimens (499 ACs). Because asymmetric PCR generates non-canonical AC morphologies, the augmentation model used for self-supervised learning was adapted accordingly. ACs were fitted using a sigmoid function complemented by a hinge linear component capturing the later single-stranded amplification phase (Section 2.9, Figure 2a), enabling the generation of realistic augmented views while preserving the characteristic morphology of the encoded signals.

We compared three approaches for classification of bulk qPCR contrived specimens (models 3.1– 3.3; Figure 3 and Figure S8). The high intrinsic separability generated by stoichiometric encoding was reflected in a baseline classification accuracy of 86.2% (chance level 12.5%), which increased to 87.4% with domain adaptation alone and to 89.8% when self-supervised pretraining was combined with domain adaptation (Figure 3). Thus, stoichiometric encoding and computational representation learning provide complementary gains: the former increases the information content and intrinsic separability of the molecular signal, while the latter enables these engineered features to be decoded across experimental domains.

**Figure 3.**
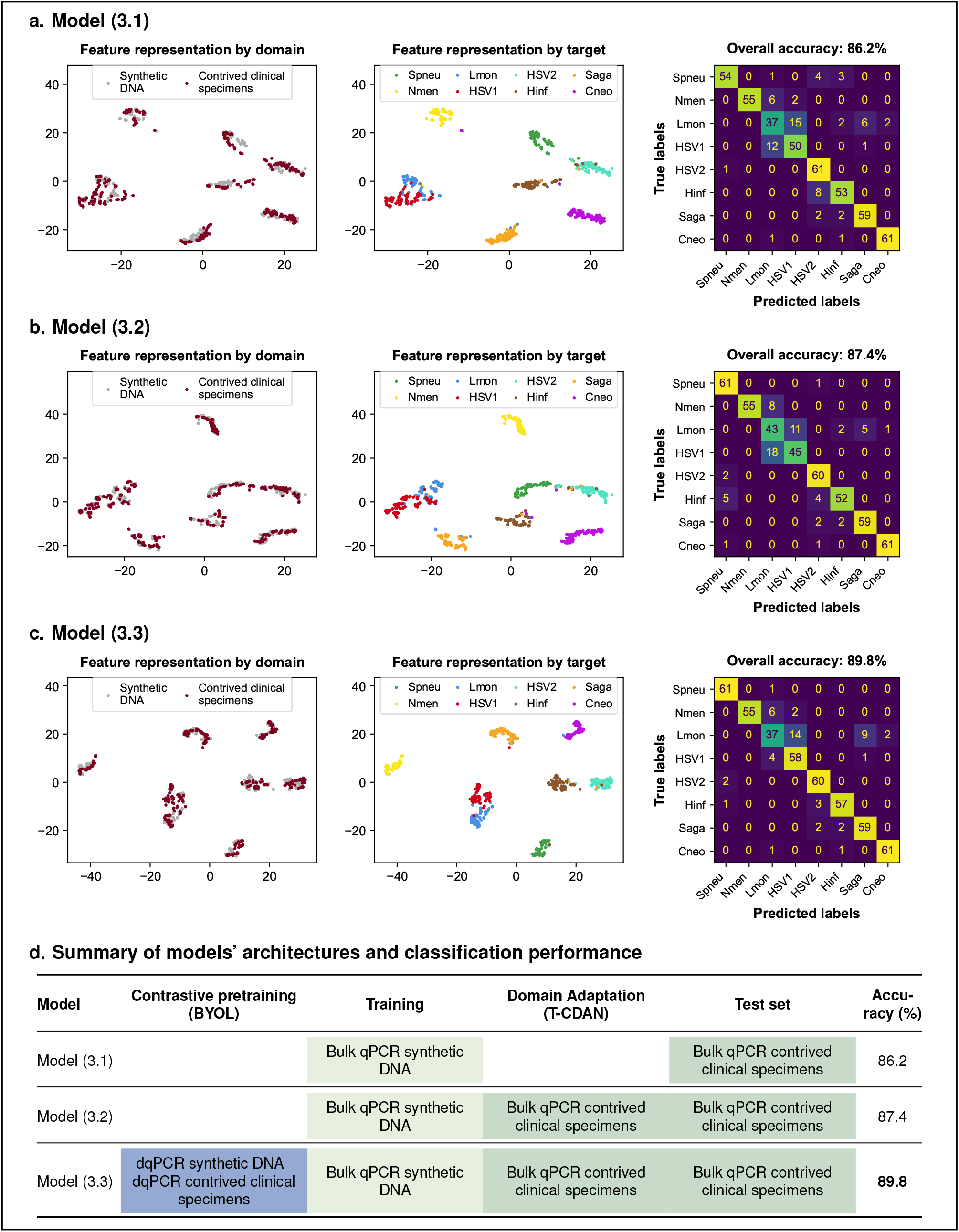
Computational decoding of stoichiometrically encoded signals in an 8-plex TaqMan assay: feature representation and classification performance. **a, b, c. t-SNE embeddings of the feature representations of bulk qPCR curves (synthetic DNA, source; contrived specimens, target) coloured by domain (left) and by class (right), and confusion matrices on the contrived specimen test set (n = 499)**. Three models were evaluated: (3.1) a 1D-transformer trained on bulk qPCR synthetic DNA data, (3.2) T-CDAN for domain adaptation between synthetic DNA (source) and contrived clinical specimens (target) data, and (3.3) T-CDAN with prior self-supervised pretraining on dqPCR data using BYOL. **d.Summary of model configurations and test accuracy**. Classification accuracy improves from 86.2% (baseline) to 87.4% with domain adaptation, and to 89.8% with the addition of self-supervised pretraining. These results demonstrate that stoichiometric feature encoding, combined with self-supervised pretraining and domain adaptation, enhances classification performance in bulk qPCR under limited labelled data conditions.

### 3.5 Stoichiometric encoding enables discrimination of co-amplification events

An important challenge for ACA is the identification of co-amplification events, in which multiple molecular targets amplify simultaneously within the same reaction. While single-target identification may be sufficient in some diagnostic settings, polymicrobial infections occur across several infectious syndromes, including respiratory and bloodstream infections (Rolston et al., 2007; Deol and Miura, 2025). A clinically useful high-plex ACA framework should therefore be able to distinguish mixed from single-target amplification profiles.

To investigate this capability under controlled conditions, we selected three targets from the encoded CNS panel: Spne, Lmon and Saga (Section 2.11). ACs were first generated for each target individually in bulk qPCR (Figure 4a, left). We then generated all three pairwise co-amplification conditions (Spne + Lmon, Spne + Saga and Lmon + Saga), together with the triple-target combination (Spne + Lmon + Saga). Mixed reactions generated reproducible composite AC morphologies that retained features of their constituent single-target profiles (Figure 4a, middle and right). The fluorescence trajectories were consistent with an approximately additive behaviour, suggesting that co-amplification generates composite kinetic signals through the superposition of contributions from the individual amplification reactions (although competition for shared reaction components and a non-linear relationship between the amount of cleaved probe and fluorescence may cause sub-additive behaviour at late cycles).

**Figure 4.**
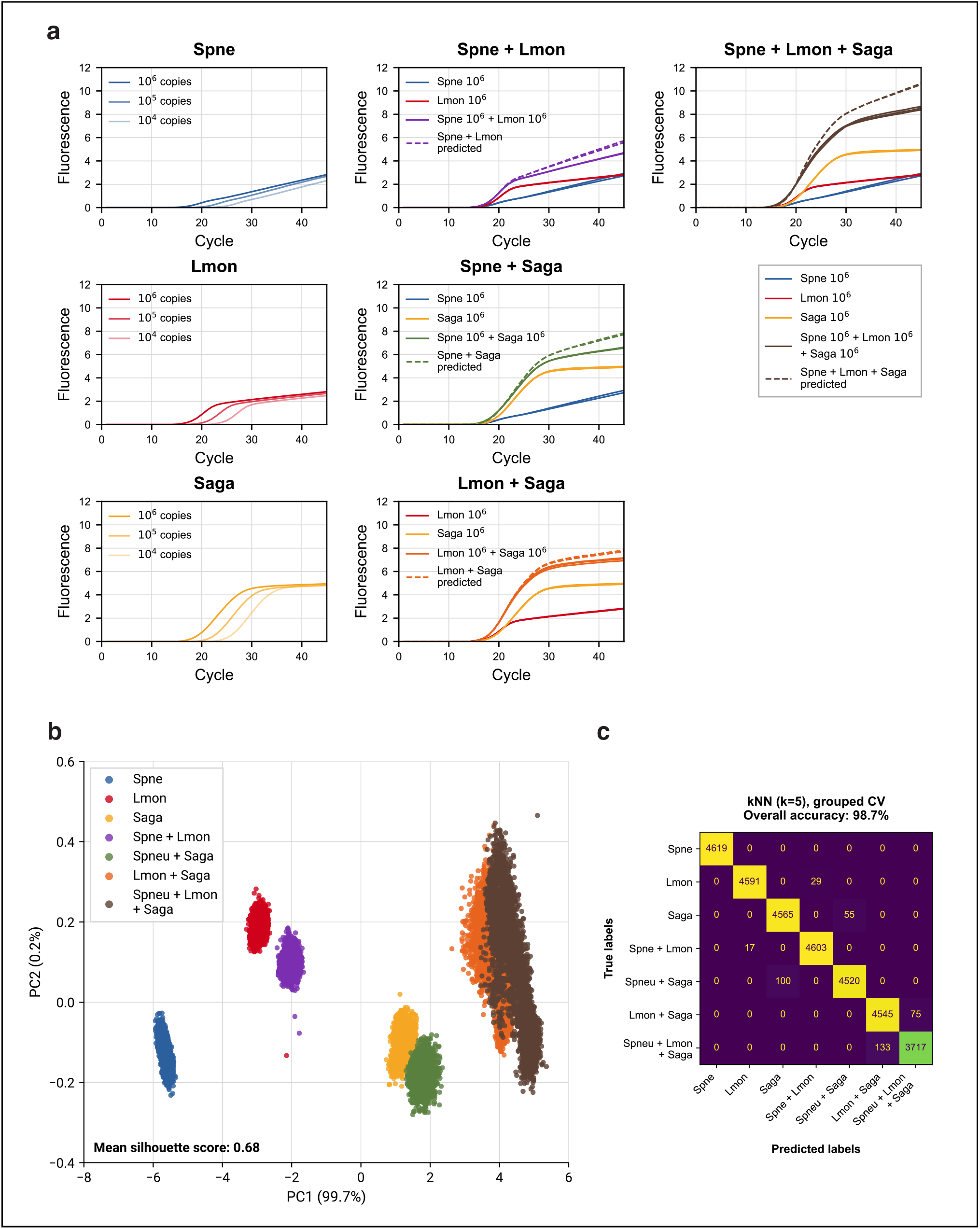
Stoichiometric encoding enables discrimination of multi-target amplification events. **a.Bulk qPCR ACs in single- and multi-target conditions**. Bulk qPCR ACs are shown for three individual encoded targets—*Streptococcus pneumoniae* (Spne), *Listeria monocytogenes* (Lmon) and *Streptococcus agalactiae* (Saga)—alone (left), in their pairwise combinations (middle) and in the triple combination (right). Single-target ACs are shown at 10^6^, 10^5^ and 10^4^ copies per reaction, co-amplification reactions are shown with equal inputs of 10^6^ copies per target (all in triplicate). Dashed lines represent the sum of fluorescence signals from the corresponding single-target reactions, while solid lines show the observed co-amplification signals. Composite kinetic profiles retain features of the contributing targets, consistent with an approximately additive fluorescence behaviour. **b.2D-PCA visualisation of co-amplification conditions**. 2D-PCA of dqPCR ACs (31,569) generated across seven conditions (three single-target, three pairwise combinations and one triple coamplification) reveals clear clustering, with a mean silhouette score of 0.68, highlighting the separability of mixed amplification profiles in feature space. **c.Classification performance**. A k-nearest neighbours classifier (*k* = 5) trained using grouped cross-validation achieves an overall accuracy of 98.7%, demonstrating accurate discrimination of both single-target and co-amplification events under controlled conditions.

We next examined whether these composite profiles remained distinguishable in partitioned reactions. Using dqPCR, we generated 31,569 ACs across seven amplification conditions comprising three single-target, three double-target and one triple-target reaction. These seven classes formed distinct clusters in 2D-PCA, with a mean silhouette score of 0.680 (Figure 4b). To quantify their separability, we trained a kNN classifier (*k* = 5) using grouped cross-validation (Section 2.11). The classifier achieved an overall accuracy of 98.7% (standard deviation [SD] across folds 0.26%]) across single-, double- and triple-target conditions (Figure 4c). These results demonstrate that stoichiometrically encoded ACs retain sufficient information to distinguish controlled multi-target amplification events with equal template inputs, extending ACA beyond single-target identification.

## 4. Discussion

In this study, we present a combined experimental and computational framework to enable high-level multiplexing in single-channel single-tube TaqMan qPCR. By integrating stoichiometric encoding with modern representation learning and adversarial domain adaptation, we demonstrate that ACs embed both thermodynamic and kinetic information that can be leveraged for diagnostic purposes, extending their role beyond their traditional use for quantification or as simple indicators of amplification success to actively engineered signals. Importantly, by leveraging signals that are already generated by standard qPCR platforms, we provide a robust, affordable, and scalable strategy to increase multiplexing capacity without requiring modifications to existing hardware or chemistry.

Building on our previous work (Moniri, Miglietta, Malpartida-Cardenas, et al., 2020; Moniri, Miglietta, Holmes, et al., 2020; Miglietta, Moniri, et al., 2021), we show that although ACs arise from a common biochemical process, they contain subtle yet reproducible signatures that can be exploited for target identification. Our results confirm that even visually similar ACs encode sufficient discriminative information to enable accurate classification using AI-based models. However, this information is inherently limited in canonical PCR settings, where ACs follow a highly constrained sigmoid morphology. To address this limitation, we introduced stoichiometric encoding as a strategy to selectively engineer non-canonical AC shapes through asymmetric PCR. This approach generates novel, distinct, and reproducible AC features—conserved from bulk qPCR to partitioned reactions—markedly increasing the intrinsic inter-target separability of the signal and facilitating downstream classification. Without the need for assay re-design or new nucleotide sequences, this method could, in prin-ciple, be adapted to existing panels by selecting candidate non-canonical designs from our 3 *×* 3 *×* 3 design space, followed by simple panel-specific optimisation.

On the computational side, a key innovation of our framework is the use of unlabelled dqPCR data for representation learning to mitigate the scarcity of labelled datasets in bulk qPCR. Unlike bulk qPCR, dqPCR readily generates large numbers of ACs from either synthetic DNA or clinical samples. We leveraged this property by designing a custom augmentation strategy that shifts ACs along the cycle axis, mimicking the variability observed in bulk qPCR due to differences in input DNA concentration, while preserving the underlying thermodynamic and kinetic molecular signatures of each target. This enabled the generation of large augmented datasets capturing realistic qPCR variability without requiring additional labelled bulk qPCR data. Self-supervised pretraining using BYOL consistently improved classification performance, indicating that the model learns robust and biologically meaningful representations of AC morphology before domain adaptation. Adversarial adaptation through T-CDAN could then align source and target domains within this structured feature space, improving transfer from synthetic to clinical bulk qPCR data. Importantly, these two components— stoichiometric feature encoding and representation learning—are complementary. While the former increases the intrinsic separability of the signal, the latter improves the ability of models to exploit this structure under realistic conditions. Together, they define a general framework in which signal generation and signal interpretation are co-designed to maximise multiplex diagnostic performance.

Our technology expands on previous work leveraging ACs to increase the number of targets per fluorescence channel in TaqMan multiplex PCR (mPCR). Amplitude modulation (Chromacode Inc), introduces controlled variations in probe concentration across targets detected within a single fluorescence channel to modulate the FFI, corresponding to the plateau of the AC (Rajagopal et al., 2019). A key advantage of this approach is its relative simplicity and its ability to disentangle coamplification events: assigning probe concentrations in powers of two (e.g., x, 2x, and 4x) ensures that each possible target combination generates a unique cumulative digested probe concentration, and therefore a unique fluorescence level. However, an intrinsic limitation of amplitude modulation is that classification relies on a single feature of the AC (the FFI). In contrast, our implementation of ACA uses the entire AC as input to AI classifiers and introduces additional non-canonical features through asymmetric PCR, effectively increasing the dimensionality of the encoded signal and enabling richer target discrimination. Another recently described approach, colour cycle multiplex amplification (CCMA), uses oligonucleotide blockers to introduce controlled delays in target amplification, thereby generating predefined permutations of fluorescence emission across multiple channels (W. Chen et al., 2024); targets are identified according to the temporal order of fluorescence emission. Nevertheless, the robustness of CCMA in the context of co-amplification events and low template concentrations remains to be fully established. Recent digital PCR approaches, binary spatial–spectral encoding (Shen et al., 2026), combinatorial barcoding (Liu et al., 2026) and FlexPCR (Park et al., 2025), achieve high multiplexing with few colours but rely on partition-level read-outs and are therefore not transferable to bulk qPCR; topology-encoded diagnostics (Lee et al., 2026) require dedicated amplicon designs. Stoichiometric encoding is, to our knowledge, the first approach to expand single-channel capacity in bulk qPCR by deliberately engineering the full temporal morphology of amplification curves using only changes in primer and probe stoichiometry on conventional thermocyclers.

Several limitations should be acknowledged. First, validation was performed on contrived specimens generated by spiking synthetic DNA into a human-DNA background derived from respiratory swabs. These do not fully capture the complexity of clinical samples, particularly in the presence of PCR inhibitors or complex microbial communities, and, for the CNS panel, do not reproduce the cerebrospinal-fluid matrix. Clinical validation using residual specimens is therefore an important next step. Second, stoichiometric encoding relies on the use of a limiting primer at low concentration, and its effect on analytical sensitivity near the limit of detection must be characterised for each target. Third, our experiments were conducted on a limited set of real-time amplification platforms, and validation across a broader range of instruments and chemistries will be required to establish general applicability. Fourth, the T-CDAN framework assumes a single source and target domain, whereas real-world applications may involve multiple, continuously evolving domains, necessitating more flexible or continual adaptation strategies. Finally, the gains conferred by domain adaptation and self-supervised pretraining were largest when domain shift was severe, as in the bulk 7-plex panel, but more modest when baseline performance was already high, as in dqPCR and the encoded 8-plex panel, suggesting that further improvements may require advances in both model architecture and data augmentation strategies.

From an experimental perspective, stoichiometric encoding introduces both opportunities and challenges. Although asymmetric PCR provides a simple and effective means of modulating AC shape without altering primer or probe sequences, increasing multiplexing levels can lead to complex interactions, including primer-dimer formation and non-specific interactions between oligonucleotides and templates or amplicons, which may distort the resulting AC shapes; assay optimisation therefore remains an iterative process as multiplexing levels increase. Furthermore, baseline-subtraction procedures embedded in commercial thermocyclers may not be fully compatible with the linear signal component characteristic of asymmetric PCR, potentially attenuating engineered kinetic features. These observations highlight the need for data-processing workflows adapted to non-canonical amplification dynamics; while we provide such workflows in this study, their implementation may require adjustments to existing analysis pipelines. Finally, the co-amplification experiments used equal template inputs in dqPCR; performance with unequal target abundances, competitive amplification dynamics and in bulk qPCR remains to be assessed.

## 5. Conclusions

Overall, this study establishes a proof-of-concept for combining stoichiometric feature engineering with modern representation learning to unlock high-level multiplexing in TaqMan qPCR-based diagnostics. By leveraging kinetic information that is already generated by standard qPCR instruments, this approach offers a practical and scalable complement to existing multiplexing strategies, without requiring additional hardware or complex assay redesign. Beyond infectious disease diagnostics, the framework may be applicable to a wide range of nucleic acid amplification technologies, including isothermal methods (Malpartida-Cardenas, Miglietta, et al., 2022) and emerging low-cost detection platforms (Malpartida-Cardenas, Moser, et al., 2023). Future work should focus on clinical validation, extension to larger multiplex panels and unequal target abundances, and evaluation across a broader range of instruments and assay conditions.

More broadly, the combination of programmable molecular systems and data-driven analysis opens new avenues for the design of diagnostic assays in which signal generation and analysis are cooptimised. This paradigm shift—from standard passive measurement to active encoding of information— provides a general design principle for next generation molecular diagnostics.

## 8. Declarations

### 8.1 Funding

This work was supported by a research scholarship from Open Innovation & Partnerships at bioMérieux (Louis Kreitmann), the Wellcome Trust funded programme CAMO-Net [226691/Z/22/Z] (Alison H. Holmes, Jesus Rodriguez-Manzano), the Imperial College President’s PhD Scholarship 2021 (Ke Xu), the Imperial College President’s PhD Scholarship 2023 (Ye Mao), and the Department of Health and Social Care-funded Center for Antimicrobial Optimisation (CAMO) at Imperial College London. Jesus Rodriguez-Manzano is supported in part by the NIHR Imperial Biomedical Research Centre (BRC).

### 8.2 CRediT authorship contribution statement

Louis Kreitmann: Conceptualization, Methodology, Investigation, Formal analysis, Visualization, Funding acquisition, Writing – original draft, review & editing. Kenny Malpartida-Cardenas: Conceptualization, Methodology, Investigation, Writing – review & editing. Ye Mao: Conceptualization, Methodology, Software, Formal analysis, Writing – review & editing. Karolina Sobala: Conceptualization, Methodology, Investigation, Writing – review & editing. Zexuan Zhao: Investigation, Writing– review & editing. San Chun Hin: Investigation, Writing – review & editing. Anirudhha Hazarika: Software, Writing – review & editing. Ke Xu: Methodology, Software, Writing – review & editing. Luca Miglietta: Methodology, Software, Writing – review & editing. Zara Breese: Investigation, Data curation, Writing – review & editing. Alison H. Holmes: Funding acquisition, Writing – review & editing. Karen Brengel-Pesce: Supervision, Funding acquisition, Writing – review & editing. Laurent Drazek: Supervision, Writing – review & editing. Jesus Rodriguez-Manzano: Conceptualization, Supervision, Project administration, Funding acquisition, Writing – review & editing.

### 8.3 Declaration of competing interests

Louis Kreitmann has received speaking fees and a research scholarship from bioMérieux, and has been employed by bioMérieux and Transgene. Jesus Rodriguez-Manzano has financial interest in ProtonDx Ltd. Alison H. Holmes is a senior investigator for the National Institute for Health Research. Karen Brengel-Pesce and Laurent Drazek are both bioMérieux employees. These authors declare that they do not have any other known competing financial interests or personal relationships that could have appeared to influence the work reported in this paper. The remaining authors declare no competing interests.

### 8.4 Data & code availability

The source data underlying all figures and the datasets required to reproduce the main analyses are available at `10.6084/m9.figshare.33685081`. Additional raw instrument files are available from the corresponding author upon reasonable request, subject to institutional and/or commercial data-sharing agreements where applicable. Custom code used for data processing, curve augmentation, model training and evaluation is available at https://github.com/louiskreitmann/ACA_BYOL_qPCR. The trained model checkpoints used for the analyses reported in this manuscript are available at 10.6084/m9.figshare.33685081.

### 8.5 Declaration of generative AI and AI-assisted technologies in the manuscript preparation process

During the preparation of this work, the authors used ChatGPT (OpenAI) and Claude (Anthropic) to assist with language editing and manuscript organisation. After using this tool, the authors reviewed and edited the content as needed and take full responsibility for the content of the publication.

### 8.6 Ethics approval

Use of residual anonymised respiratory swabs was approved under IRAS Project ID 289096. All samples were anonymised before use.

## Notes

https://github.com/louiskreitmann/ACA_BYOL_qPCR

https://figshare.com/articles/dataset/ACA_BYOL_qPCR_datasets_and_model_checkpoints/33685081

### Competing Interest Statement

Louis Kreitmann has received speaking fees and a research scholarship from bioMerieux, and has been employed by bioMerieux and Transgene. Jesus Rodriguez-Manzano has financial interest in ProtonDx Ltd. Alison H. Holmes is a senior investigator for the National Institute for Health Research. Karen Brengel-Pesce and Laurent Drazek are both bioMerieux employees. These authors declare that they do not have any other known competing financial interests or personal relationships that could have appeared to influence the work reported in this paper. The remaining authors declare no competing interests.

