## Supplementary Material for "Stoichiometric Encoding of Amplification Kinetics Enables High-Level Multiplexing in Single-Channel TaqMan Real-Time PCR"

### ELECTRONIC SUPPLEMENTARY MATERIAL


#### Authors

Louis Kreitmann[1,2,†], Kenny Malpartida-Cardenas[1,†], Ye Mao[3,†], Karolina Sobala[1], Zexuan Zhao[1], San Chun Hin[1], Anirudhha Hazarika[1], Ke Xu[1,3], Luca Miglietta[1], Zara Breese[1], Alison H. Holmes[1,4], Karen Brengel-Pesce[5], Laurent Drazek[6] and Jesus Rodriguez-Manzano[1,4,*]

#### Affiliations

[1]Department of Infectious Disease, Imperial College London, London, UK
[2]Open Innovation & Partnerships, bioMérieux, Marcy-l'Étoile, France
[3]Department of Electrical and Electronic Engineering, Imperial College London, London, UK
[4]The Fleming Initiative, Imperial College London and Imperial College Healthcare NHS Trust, London, UK
[5]Molecular Biology, Research & Development, bioMérieux, Grenoble, France
[6]Data Science, Research & Development, bioMérieux, Grenoble, France
[†]LK, KMC, and YM contributed equally to this work as first authors.

### SUPPLEMENTARY METHODS

#### Primers and assay design

Primer design and *in silico* analysis to evaluate the presence of secondary structures (e.g., hairpins), primer dimers and self-dimers were performed using Geneious Prime 2023.0 (Biomatters, New Zealand). Off-target binding was assessed using the embedded Geneious BLAST tool. Key primer characteristics, including melting temperature ($T_m$) and GC content, were extracted from Geneious Prime. Further analysis to avoid primer dimers and secondary structures was performed using the Multiple Primer software (ThermoFisher, United States) and Nupack.

All primers were synthesised by Integrated DNA Technologies (IDT, United States). TaqMan probes were double-quenched hydrolysis probes labelled with 6-FAM as the fluorophore at the 5′ end, a ZEN internal quencher, and a 3′ Iowa Black® FQ (3IABkFQ) quencher, also purchased from IDT. In all double-quenched probes, the ZEN moiety was located between the 9$^{th}$ and the 10$^{th}$ nucleotide (in the 5′ to 3′ direction). All oligos were resuspended in TE buffer at a concentration of 100 $\mu$mol $\cdot$ L$^{-1}$.

#### Synthetic DNA templates

Synthetic DNA fragments (gBlocks) were purchased from IDT, resuspended in TE buffer to a concentration of 5 ng/$\mu$L, and stored at $-20$°C. DNA concentrations were quantified using a NanoDrop One spectrophotometer (ThermoFisher, United States), and 10-fold serial dilutions were prepared in TE buffer.

#### Bulk qPCR experiments

All experiments were conducted on a QuantStudio™ 5 Real-Time PCR System (ThermoFisher, United States). qPCR mixes contained $5$ $\mu$L 2X IDT PrimeTime Gene Expression Master Mix with low-concentration ROX (catalogue number 1055772), $1$ $\mu$L 20X GE, $1$ $\mu$L 10X primer mix, and $1$ $\mu$L DNA templates (or NTCs). The final reaction volume was adjusted to $10$ $\mu$L with nuclease-free water. Thermal cycling conditions consisted of a hot-start step at $95\,°\mathrm{C}$ for 3 minutes, followed by 50-60 cycles at $95\,°\mathrm{C}$ for 15 seconds (denaturation) and $60\,°\mathrm{C}$ for 30-45 seconds (annealing and elongation).

#### dqPCR experiments

All experiments were conducted on a HD/Biomark (Fluidigm Corporation, South San Francisco, California, United States) using the qdPCR 37K™ integrated fluidic circuit (IFC) provided by the same company. The priming and loading steps of the IFC were followed as per the supplier's protocol (Fluidigm document number 100-6896 Rev 03). dqPCR mixes contained $3$ $\mu$L 2X IDT PrimeTime Gene Expression Master Mix (with passive ROX), $0.6$ $\mu$L 20X GE, $0.6$ $\mu$L 10X primer mix, $1.8$ $\mu$L DNA templates (or NTCs). The final reaction volume was adjusted to $6$ $\mu$L with nuclease-free water. A total of $4.5$ $\mu$L of reaction mix was transferred to each inlet (or panel) of a Fluidigm 37K IFC for the thermal cycling step. PCR cycling conditions followed those used for bulk qPCR experiments.

### DNA extraction from nasal swabs

Residual and anonymised respiratory swabs were obtained through the virology laboratory at Imperial College Healthcare NHS Trust (IRAS Project ID 289096). All samples had previously tested negative for an extended panel of respiratory viruses using conventional diagnostic methods. Samples were extracted using the QIAsymphony® DSP Virus/Pathogen Midi Kit (Cat. No. 937055, QIAGEN) in combination with the QIAsymphony SP instrument. Briefly, the Complex 400 protocol was used, processing $400\ \mu L$ of input sample. Elution was performed in cooled 96-well low-skirt plates with a final elution volume of $110\ \mu L$ per sample. Eluates were tested on the same day and subsequently stored at $-80\ °C$.

### 7-plex TaqMan panel for respiratory viruses

DNA templates for the 7-plex TaqMan panel for respiratory viruses are presented in Table S1.

| Microorganism | Abbreviation | Target gene (abbreviation) | NCBI Accession No. | Template length (bp) | Amplicon length (bp) |
|---|---|---|---|---|---|
| HAdV (Adenovirus) | Adeno | Hexon (HEX) | NC_001405 | 300 | 129 |
| SARS-CoV-2 | COVID | Nucleocapsid (N) | NC_045512 | 200 | 78 |
| HCoV-229E | C22 | Nucleocapsid (N) | NC_002645 | 570 | 158 |
| HCoV-HKU1 | CHK | Nucleocapsid (N) | NC_006577 | 300 | 206 |
| HCoV-NL63 | CNL | Nucleocapsid (N) | NC_005831 | 300 | 77 |
| HCoV-OC43 | COC | Nucleocapsid (N) | NC_006213 | 300 | 115 |
| MERS-CoV | MERS | Nucleocapsid (N) | NC_019843 | 300 | 67 |

**Table S1: Synthetic DNA templates for the 7-plex TaqMan panel for respiratory viruses.**

Primers and probes for the 7-plex TaqMan panel for respiratory viruses are presented in Table S2.

| **Microorganism (abbreviation)** | **Target gene (abbreviation)** | **Oligo type** | **Oligo sequence (5′-3′)** | **Oligo $T_m$ (°C)** |
|---|---|---|---|---|
| HAdV (Adeno) | Hexon (HEX) | forward | CGCAGTGGTCTTACATGCACATCTC | 69.72 |
| | | reverse | GCCACCGTGGGGTTTCTAAACTTG | 70.15 |
| | | probe | CCTCGGAGTACCTRAGCCCCGG | 72.94 |
| HCoV-229E (C22) | Nucleocapsid (N) | forward | GAAATGCAAAAGCCACGGTGGAA | 68.67 |
| | | reverse | AGCTCAGCAAATTGTGGATAGCC | 67.37 |
| | | probe | AGTTGTGGTCAAGGTCTCTGGGGCC | 71.21 |
| HCoV-HKU1 (CHK) | Nucleocapsid (N) | forward | TCAAGAAGCTATCCCTACTAGGT | 63.71 |
| | | reverse | GCGATCTCATCAGCCATATCAGGT | 68.29 |
| | | probe | CGCCTGGTACGATTTTGCCTCAAGGCT | 72.21 |
| HCoV-NL63 (CNL) | Nucleocapsid (N) | forward | GGTGCTAAAACTGTTAATACCAGT | 62.95 |
| | | reverse | GCAATAGAGAACTTTGGTTCCA | 62.40 |
| | | probe | AGGTTTCTGATTACGTTTGCGATTACCA | 66.42 |
| HCoV-OC43 (COC) | Nucleocapsid (N) | forward | GGTGGAGAAATGTTAAAACTTGGAACT | 66.26 |
| | | reverse | CCAAAGAAAAACGCACCAGCTG | 66.94 |
| | | probe | TCCCCATTCTTGCAGAACTCGCACCCA | 73.13 |
| SARS-CoV-2 (COVID) | Nucleocapsid (N) | forward | ATAATGGACCCCAAAATCAGCGA | 66.41 |
| | | reverse | TCTGGTTACTGCCAGTTGAATCTG | 66.66 |
| | | probe | CACCCCGCATTACGTTTGGTGGACC | 70.86 |
| MERS-CoV (MERS) | Nucleocapsid (N) | forward | ACGCGGAACCCTAACAATGATT | 66.51 |
| | | reverse | GTGGGTCCTCAGTGCCGAGT | 69.98 |
| | | probe | TTCCCTGGAGGTCTCCTGGTCCGC | 72.33 |

**Table S2: Primers and probes for the 7-plex TaqMan panel for respiratory viruses.**
TaqMan probes were double-quenched hydrolysis probes labelled with 6-FAM as the fluorophore at the 5′ end, a ZEN internal quencher, and a 3′ Iowa Black® FQ (3IABkFQ) quencher, also purchased from IDT. In all double-quenched probes, the ZEN moiety was located between the 9$^{th}$ and the 10$^{th}$ nucleotide (in the 5′ to 3′ direction). For instance for the Adenovirus target, the exact sequence was 5′-/56-FAM/CCTCGGAGT/ZEN/ACCTRAGCCCCGG/3IABkFQ/-3′.
$T_m$: melting temperature.

### 8-plex TaqMan panel for central nervous system infections

DNA templates for the 8-plex TaqMan panel for CNS infections are presented in Table S3.

| Microorganism | Abbreviation | Target gene (abbreviation) | NCBI Accession No. | Template length (bp) | Amplicon length (bp) |
|---|---|---|---|---|---|
| *Streptococcus pneumoniae* | Spne | Pneumolysin (*ply*) | M17717.1 | 1003 | 81 |
| *Neisseria meningitidis* | Nmen | Capsule transport protein A (*ctrA*) | GU391295 | 1018 | 111 |
| *Listeria monocytogenes* | Lmon | Listeriolysin O (*hly*) | X61210 | 1002 | 112 |
| Herpes simplex virus 1 | HSV1 | Glycoprotein D (gD) | JQ673480.1 | 1283 | 71 |
| Herpes simplex virus 2 | HSV2 | Glycoprotein G (gG) | AJ303204 | 1009 | 111 |
| *Haemophilus influenzae* | Hinf | Capsule export protein A (*bexA*) | L42023 | 1007 | 108 |
| *Streptococcus agalactiae* | Saga | CAMP factor (*cfb*) | X72754.1 | 936 | 93 |
| *Cryptococcus neoformans* | Cneo | 18S ribosomal DNA (18S rDNA) | AF075483.1 | 1033 | 124 |

**Table S3: DNA templates for the 8-plex TaqMan panel for CNS infections.**

Primers and probes for the 8-plex TaqMan panel for CNS infections, along with concentrations used for feature encoding, are presented in Table S4.

| **Micro-organism abbreviation** | **Target gene (abbreviation)** | **Oligo type** | **Sequence** | **Oligo** $T_m$ | **Concentration (nmol·L$^{-1}$)** |
|---|---|---|---|---|---|
| Spne | Pneumolysin (*ply*) | forward | TGCAGAGCGTCCTTTGGTCTAT | 60.3 | 150 |
| | | reverse | CTCTTACTCGTGGTTTCCAACTTGA | 61.3 | 150 |
| | | probe | TGGCGCCCATAAGCAACACTCGAA | 64.4 | 150 |
| Nmen | Capsule transport protein A (*ctrA*) | forward | GCTGCGGTAGGTGGTTCAA | 58.8 | 40 |
| | | reverse | TTGTCGCGGATTTGCAACTA | 55.3 | 450 |
| | | probe | CATTGCCACGTGTCAGCTGCACAT | 64.4 | 450 |
| Lmon | Listeriolysin O (*hly*) | forward | TGCAAGTCCTAAGACGCCA | 56.7 | 450 |
| | | reverse | CACTGCATCTCCGTGGTATACTAA | 61.0 | 450 |
| | | probe | CGATTTCATCCGCGTGTTTCTTTTCG | 63.2 | 80 |
| HSV1 | Glycoprotein D (gD) | forward | CGGCCGTGTGACACTATCG | 61.0 | 450 |
| | | reverse | CTCGTAAAATGGCCCCTCC | 58.8 | 450 |
| | | probe | CCATACCGACCACACCGACGAACC | 67.8 | 450 |
| HSV2 | Glycoprotein G (gG) | forward | CGCTCTCGTAAATGCTTCCCT | 59.8 | 450 |
| | | reverse | TCTACCCACAACAGACCCACG | 61.8 | 450 |
| | | probe | CGCGGAGACATTCGAGTACCAGATCG | 68.0 | 150 |
| Hinf | Capsule export protein A (*bexA*) | forward | TGCGGTAGTGTTAGAAAATGGTATTATG | 60.7 | 450 |
| | | reverse | GGACAAACATCACAAGCGGTTA | 58.4 | 150 |
| | | probe | ACAAAGCGTATCAATACTACAAC<br>GAGACGCAAAAA | 66.0 | 450 |
| Saga | CAMP factor (*cfb*) | forward | AGGGAACAGATTATGAAAAACCGG | 59.3 | 150 |
| | | reverse | AAGGCTTCTACACGACTACCAA | 58.4 | 450 |
| | | probe | AGACTTCATTGCGTGCCAACCCTGAGAC | 68.0 | 450 |
| Cneo | 18S ribosomal DNA (18S rDNA) | forward | TCCTCACGGAGTGCACTGTCTTG | 64.2 | 450 |
| | | reverse | CAGTTGTTGGTCTTCCGTCAATCTA | 61.3 | 300 |
| | | probe | TCCTGGTTCCCCTGCACACCCA | 65.8 | 450 |

**Table S4: Primer and probe sequences and concentrations in the 8-plex TaqMan panel for CNS infections.**
TaqMan probes were double-quenched hydrolysis probes labelled with 6-FAM as the fluorophore at the 5′ end, a ZEN internal quencher, and a 3′ Iowa Black® FQ (3IABkFQ) quencher, also purchased from IDT. In all double-quenched probes, the ZEN moiety was located between the 9$^{th}$ and 10$^{th}$ nucleotides (in the 5′ to 3′ direction). For instance, for the Spne target, the exact probe sequence was 5′-/56-FAM/TGGCGCCCA/ZEN/TAAGCAACACTCGAA/3IABkFQ/-3′. All concentrations are expressed in nmol·L$^{-1}$.

**Pre-processing of amplification data: detailed procedure**

ACs obtained in dqPCR and qPCR undergo a series of post-processing steps before being used as input data to ML models. These include:

- *Removing 'no amplification' data points*. In both bulk qPCR and dqPCR experiments, we include reaction wells for NTCs, containing the PCR mix (dNTPs, Taq polymerase, buffer, primers and probes) but no template; these wells show no amplification (flat fluorescent signal) and are removed from the set of training and testing data. Second, in dqPCR when the initial concentration of template is lower than the threshold of saturation of the 37K digital chip, some of the wells in the chip also show no amplification, and these data points are also discarded.
- *Baseline correction*. Because of background fluorescence, the initial flat portion of ACs is not always set at a null level of fluorescence. To ensure that the baseline of all curves is zero, we apply an in-house cycle threshold ($C_t$)-based correction algorithm. For each AC, we compute the second derivative of the fluorescence signal to identify the $C_t$ (defined as the cycle with the maximum second derivative of fluorescence with respect to cycle number). We then calculate the mean background fluorescence over a window of $n$ cycles (with $n = 5$ to $n = 10$) immediately preceding (but excluding) the $C_t$. This mean value is subtracted from the raw fluorescence data of the entire curve, and any resulting negative fluorescence values are floored to zero.
- *Inter-experiment normalisation*. This step is crucial because: (1) data obtained on different thermocyclers (or even from different experiments run on the same thermocycler) can have substantially different scales (i.e., the fluorescence values obtained at the last cycle, corresponding to the plateau of ACs, are different for a given assay); and (2) the classification accuracy of deep learning models for ACA depends on the relative scale of ACs obtained in any $n$-plex panel of interest. To ensure that different datasets remain on a comparable scale, we include in all experiments as a positive control a synthetic DNA template used for normalisation. We calculate the mean final fluorescence intensity of ACs obtained for this control (each averaged over the last $n$ cycles, usually $n = 5$), and divide the fluorescence values obtained for all curves within that 96-well plate or 37K digital chip by this value.
- *Smoothing*. To remove small experimental variations, we apply a Savitzky-Golay filter to each curve (with filter window length of 15 and order of the polynomial used to fit the samples of 3).

**T-CDAN: architecture and loss functions**

ACs are fed into the feature extractor as $c$-dimensional vectors, $c$ being the number of PCR cycles. Let $\mathbf{x} \in \mathbb{R}^c$ be an AC. $\mathbf{x}$ is standardised (mean $0$ and standard deviation $1$) and mapped to a $N_d$-dimensional latent space using a learnable token embedding layer, giving $\mathbf{E}_{\text{embed}} \in \mathbb{R}^{c \times N_d}$. A position embedding $\mathbf{E}_{\text{pos}}$ is then added element-wise to the linear projection $\mathbf{E}_{\text{embed}}$ before the resulting sum $\mathbf{Z}$ is fed into a transformer encoder. Each transformer encoder is a stack of $N$ transformer blocks, each composed of a multi-head self-attention (MHA) and a position-wise multilayer perceptron (MLP). Each encoder block outputs a representation matrix of dimension $\mathbb{R}^{c \times N_d}$, consistent with the dimension of the input embedding matrix $\mathbf{Z}$. The last embedding matrix $\mathbf{Z}_N$ is then reshaped through a simple MLP into a feature representation $\mathbf{f}$.

Mathematically, this can be expressed as:

$$\mathbf{Z}_0 = \mathbf{E}_{\text{embed}} \oplus \mathbf{E}_{\text{pos}} \tag{1}$$

$$\mathbf{Z}'_i = \mathrm{MHA}\left(\mathrm{LN}\left(\mathbf{Z}_{i-1}\right)\right) + \mathbf{Z}_{i-1}, \quad \forall\, i = 1, 2, \ldots, N \tag{2}$$

$$\mathbf{Z}_i = \mathrm{MLP}\left(\mathrm{LN}\left(\mathbf{Z}'_i\right)\right) + \mathbf{Z}'_i, \quad \forall\, i = 1, 2, \ldots, N \tag{3}$$

$$\mathbf{f} = \mathrm{MLP}\left(\mathrm{flatten}\left(\mathbf{Z}_N\right)\right) \tag{4}$$

where $\oplus$ denotes element-wise addition, $\mathbf{Z}'_i \in \mathbb{R}^{c \times N_d}$ is the output of the $i^{\text{th}}$ MHA block, $\mathbf{Z}_i \in \mathbb{R}^{c \times N_d}$ is the output of the $i^{\text{th}}$ point-wise MLP block, and $\mathbf{Z}_N$ is the output of the final transformer encoder layer.

The label predictor $G$ takes $\mathbf{f}$ as input and maps it through a fully connected layer into a vector $\mathbf{g} = G(\mathbf{f})$ of dimension $\mathbb{R}^{n_c}$, where $n_c$ is the number of classes to predict (i.e., the number of targets in the multiplex panel). Then, $\hat{\mathbf{y}} = \mathrm{softmax}(\mathbf{g})$ represents a probability distribution over the $n_c$ label classes, and $\mathcal{L}_G$, the loss function of $G$, is a categorical cross-entropy loss (negative log-likelihood):

$$\mathcal{L}_G(\theta_F, \theta_G) = -\mathbb{E}_{(\mathbf{x}^s, y^s)\sim\mathcal{S}} \sum_{i=1}^{n_c} \mathbf{1}_{y^s=i} \log(\hat{y}_i^s) \tag{5}$$

where $\theta_F$ and $\theta_G$ are the parameters of $F$ and $G$, respectively; $\mathbb{E}_{(\mathbf{x}^s, y^s)\sim\mathcal{S}}$ denotes the expectation over all training examples from the source domain $\mathcal{S}$ (each consisting of input data $\mathbf{x}^s$ and label $y^s$), $i = 1, \ldots, n_c$ indexes the class labels, and $\mathbf{1}_{y^s=i}$ is 1 if the true label $y^s$ is equal to the current class index $i$, and 0 otherwise.

The aim of the conditional domain classifier $D$ is to distinguish whether extracted features $\mathbf{f}$ from training examples $\mathbf{x}^{\mathcal{S}}$ and $\mathbf{x}^{\mathcal{T}}$ belong to either the source domain $\mathcal{S}$ or the target domain $\mathcal{T}$. The loss function of $D$, $\mathcal{L}_D$, is inspired by a cross-entropy loss, except that it does not directly take $\mathbf{f}$ as input, but rather the outer product of $\mathbf{f}$ and $\mathbf{g}$:

$$\mathbf{h} = \mathbf{f} \otimes \mathbf{g} \tag{6}$$

$$\mathcal{L}_D\left(\theta_F, \theta_D\right) = -\, \mathbb{E}_{\mathbf{x}^s \sim S} \log(D\left(\mathbf{h}^s\right)) -\, \mathbb{E}_{\mathbf{x}^t \sim T} \log(1 - D\left(\mathbf{h}^t\right)) \tag{7}$$

where $\otimes$ is the outer product, $\theta_F$ are the parameters of $F$, $\theta_D$ those of $D$, $\mathbb{E}_{\mathbf{x}^s \sim S}$ indicates the expectation over all training examples in the source domain $\mathcal{S}$ (note that labels are not provided), $\mathbb{E}_{\mathbf{x}^t \sim T}$ the expectation over all training examples in the target domain $\mathcal{T}$, and $D(\mathbf{h})$ indicates the probability of predicting $\mathbf{h}$ to belong to the source domain. From equation 7, it can be seen that $\mathcal{L}_D$ is minimised when $D\left(\mathbf{h}^s\right)$ tends to one (correctly predicting that $\mathbf{h}^s$ belongs to $\mathcal{S}$) and $D\left(\mathbf{h}^t\right)$ tends to zero (correctly predicting that $\mathbf{h}^t$ belongs to $\mathcal{T}$), while it is maximised in the opposite cases.

The goal of training is to minimise $\mathcal{L}_G$ while maximising $\mathcal{L}_D$, i.e., to find the optimal parameters $(\theta_F,\ \theta_G,\ \theta_D)$ that minimise the overall loss:

$$\mathcal{L}\left(\theta_F,\ \theta_G, \theta_D\right) = \mathcal{L}_G\left(\theta_F,\ \theta_G\right) - \lambda \mathcal{L}_D\left(\theta_F,\ \theta_D\right), \tag{8}$$

where $0 \leq \lambda \leq 1$ is a parameter for balancing the two objectives. The parameters $\theta_F$ and $\theta_G$ are updated to minimise the overall loss $L$, effectively training the feature extractor $F$ to produce features that are both discriminative for the source task (helping $G$ minimise $\mathcal{L}_G$) and confusing for the domain classifier $D$ (by aiming to maximise $\mathcal{L}_D$ through the $-\lambda\mathcal{L}_D$ term):

$$(\hat{\theta}_F, \hat{\theta}_G) = \arg\min_{\theta_F, \theta_G} \left[\mathcal{L}_G(\theta_F, \theta_G) - \lambda\mathcal{L}_D(\theta_F, \hat{\theta}_D)\right] \tag{9}$$

Concurrently, or in an alternating step, the parameters $\theta_D$ of the domain classifier $D$ are updated while keeping the parameters of the feature extractor $\hat{\theta}_F$ and label predictor $\hat{\theta}_G$ fixed. The objective for the domain classifier is to accurately differentiate between features derived from the source domain and those from the target domain. Therefore, $D$ is trained by minimising its own domain classification loss $\mathcal{L}_D$ with respect to its parameters $\theta_D$:

$$\begin{aligned} \hat{\theta}_D &= \arg\min_{\theta_D} \mathcal{L}_D(\hat{\theta}_F, \theta_D) \\ &= \arg\max_{\theta_D} -\lambda\mathcal{L}_D\left(\hat{\theta}_F,\ \theta_D\right) \\ &= \arg\max_{\theta_D} L(\hat{\theta}_F, \hat{\theta}_G, \theta_D) \end{aligned} \tag{10}$$

This process sharpens the domain classifier's ability to detect the origin of the features, setting up the adversarial challenge for the subsequent update of the feature extractor. Ultimately, this yields adversarial learning between the feature extractor/label predictor and the domain classifier, i.e., the features extracted by $F$ maximise the ability of $G$ to correctly identify class labels, while minimising the ability of $D$ to discriminate between domains.

**Algorithm 1:** Training procedure of T-CDAN

**Input :**
- Labelled ACs from synthetic DNA (source domain) $\mathcal{S} = \{(\mathbf{x}_i^s, y_i^s)\}_{i=1}^{n_s}$,
- Unlabelled ACs from contrived clinical specimens (target domain) $\mathcal{T} = \left\{\mathbf{x}_j^t\right\}_{j=1}^{n_t}$,
- Source domain label set $\mathcal{D}_s = \{1\}$ and target domain label set $\mathcal{D}_t = \{0\}$,
- Parameter $\lambda$ for weighting loss and learning rate $\mu$;

**Output:** Trained parameters $\hat{\theta}_F, \hat{\theta}_G, \hat{\theta}_D$;

Initialize model parameters and learning rate $\mu$.
**while** *pre-defined maximum iterations is not met* **do**
  Compute label prediction loss $\mathcal{L}_G(\theta_F, \theta_G)$
  Compute domain classifier loss $\mathcal{L}_D(\theta_F, \theta_D)$
  `// Update feature generator parameters`
  $\theta_F \leftarrow \theta_F - \mu(\frac{\partial\mathcal{L}_G}{\partial\theta_F} - \lambda\frac{\partial\mathcal{L}_D}{\partial\theta_F})$
  `// Update label predictor parameters`
  $\theta_G \leftarrow \theta_G - \mu\frac{\partial\mathcal{L}_G}{\partial\theta_G}$
  `// Update conditional domain classifier parameters`
  $\theta_D \leftarrow \theta_D - \mu\lambda\frac{\partial\mathcal{L}_D}{\partial\theta_D}$

### BYOL: loss function

From the first augmented view $\mathbf{v}$, the online network outputs a representation $\mathbf{y}_\theta = f_\theta(\mathbf{v})$ and a projection $\mathbf{z}_\theta = g_\theta(\mathbf{y}_\theta)$. The target network outputs $\mathbf{y'}_\xi = f_\xi(\mathbf{v'})$ and a target projection $\mathbf{z'}_\xi = g_\xi(\mathbf{y'}_\xi)$. The last part of the online network then outputs a prediction $\mathbf{q}_\theta(\mathbf{z}_\theta)$ of $\mathbf{z'}_\xi$ and computes a mean squared error between the $\ell_2$-normalised predictions and target projections:

$$\overline{\mathbf{q}}_\theta(\mathbf{z}_\theta) = \frac{\mathbf{q}_\theta(\mathbf{z}_\theta)}{\|\mathbf{q}_\theta(\mathbf{z}_\theta)\|_2} \tag{11}$$

$$\overline{\mathbf{z}}'_\xi = \frac{\mathbf{z'}_\xi}{\|\mathbf{z'}_\xi\|_2} \tag{12}$$

$$\mathcal{L}_{\theta,\xi} = \left\|\overline{\mathbf{q}}_\theta\left(\mathbf{z}_\theta\right) - \overline{\mathbf{z}}'_\xi\right\|_2^2 \tag{13}$$

This loss is symmetrised by feeding $\mathbf{v}$ into the target network and $\mathbf{v'}$ into the online network to compute $\widetilde{L}_{\theta,\xi}$. $\mathcal{L}^{BYOL}_{\theta,\xi}= \mathcal{L}_{\theta,\xi} + \widetilde{\mathcal{L}}_{\theta,\xi}$ is minimised with respect to $\theta$ but not $\xi$ :

$$\theta \leftarrow \text{optimizer}\left(\theta,\ \nabla_\theta \mathcal{L}^{BYOL}_{\theta,\xi}, \eta\right), \tag{14}$$

where $\nabla_\theta$ is the gradient with respect to $\theta$ and $\eta$ is the learning rate.

### Deep learning pipeline: implementation details

All experiments were run on an NVIDIA RTX 4090 GPU.

T-CDAN used a Transformer feature extractor for ACs, with a 128-dimensional input embedding, four Transformer blocks, four self-attention heads per block, regular positional encoding with period 20, and a 256-dimensional bottleneck. The label predictor and domain classifier were shallow MLP heads for class prediction and source/target domain prediction, respectively. Models were trained for 10,000 iterations using AdamW with a cosine learning-rate schedule. For the 8-plex CNS panel experiments, we used a learning rate of $10^{-4}$, batch size of 96, and weight decay of $10^{-6}$; for the 7-plex experiments, we used a learning rate of $2 \times 10^{-5}$, batch size of 16, and weight decay of $10^{-4}$.

The BYOL online network consisted of the same Transformer-based encoder and an MLP predictor with a hidden dimension of 4096 and output feature dimension of 256. The target network was maintained as an exponential moving average (EMA) of the online encoder, with a moving-average decay of 0.99. BYOL training used AdamW with a learning rate of $2 \times 10^{-4}$, weight decay of $10^{-4}$, $\beta = (0.9, 0.999)$, 100 warmup steps, and cosine decay to a minimum learning-rate scale of 0.02. The self-supervised dataset was split into 90% training and 10% validation. BYOL was trained with a batch size of 1024 for 100 epochs in both the 7-plex respiratory viruses and 8-plex CNS panels settings.

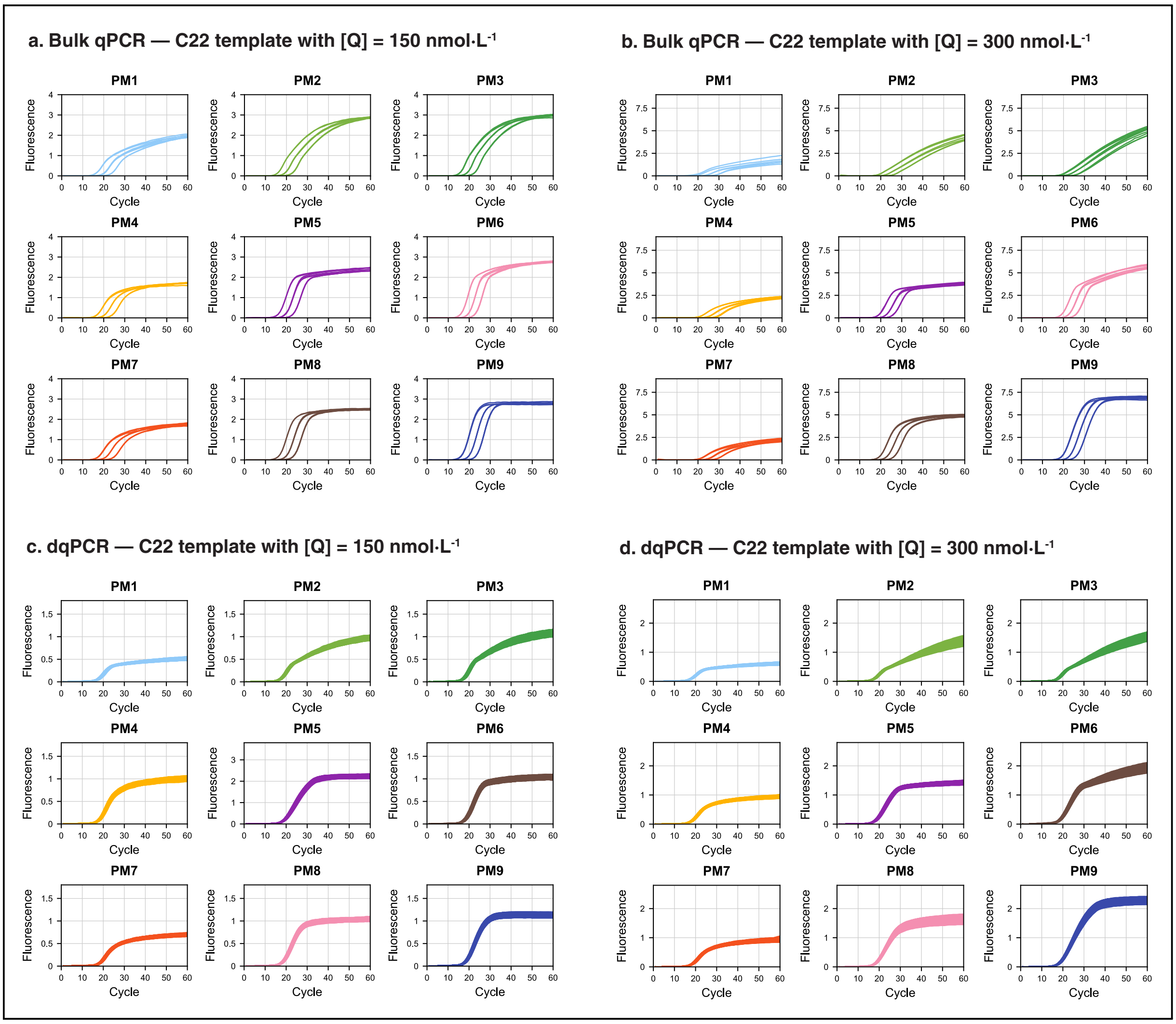


**Figure S1. Expansion of the stoichiometric design space across primer and probe concentrations in bulk and digital qPCR.**

ACs obtained for the HCoV-229E template across nine primer mixes (PM1–PM9) at two probe concentrations in bulk and digital qPCR. (a,b) Bulk qPCR experiments were performed at $[Q] = 150$ and $[Q] = 300\ \mathrm{nmol{\cdot}L^{-1}}$, respectively. (c,d) Corresponding dqPCR experiments were conducted at the same probe concentrations. Please note the variation in the y-axis scale across experiments. Bulk qPCR experiments were performed using three decreasing concentrations of input DNA ($10^6$, $10^5$, and $10^4$ copies + NTCs) in triplicate. dqPCR experiments were performed at input concentrations of $10^6$, $10^5$, and $10^4$ copies + NTCs in triplicate.

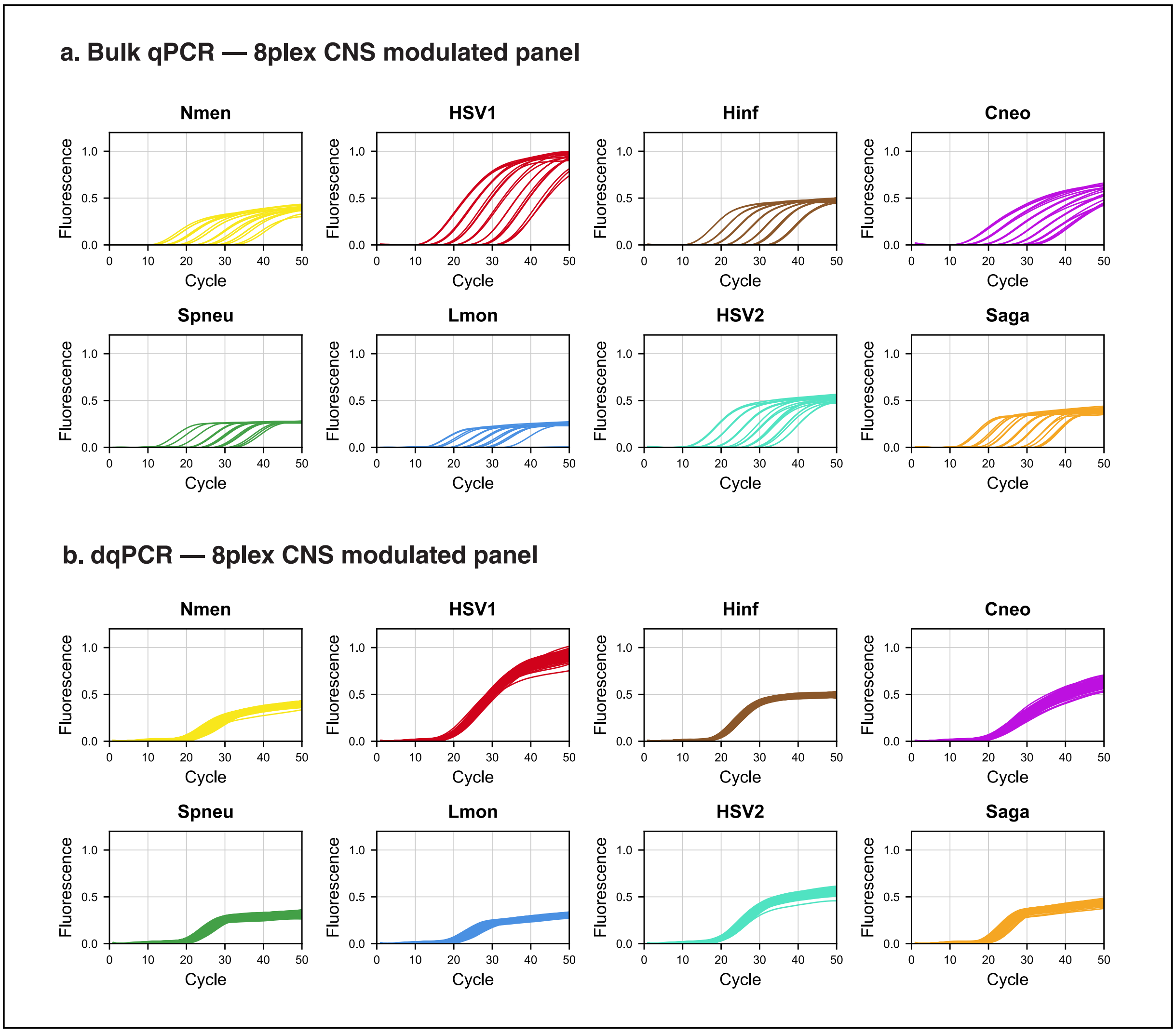


**Figure S2. Stoichiometrically encoded target-specific ACs in the 8-plex CNS infection panel.** ACs obtained for the encoded 8-plex CNS infections panel across two experimental platforms: (a) bulk qPCR (489 ACs) and (b) dqPCR (81,007 ACs). Bulk qPCR experiments were performed using decreasing concentrations of input DNA from $10^7$ copies to 1 copy + NTCs in triplicate. dqPCR experiments were performed at input concentrations of $10^6$, $10^5$, and $10^4$ copies + NTCs in triplicate.

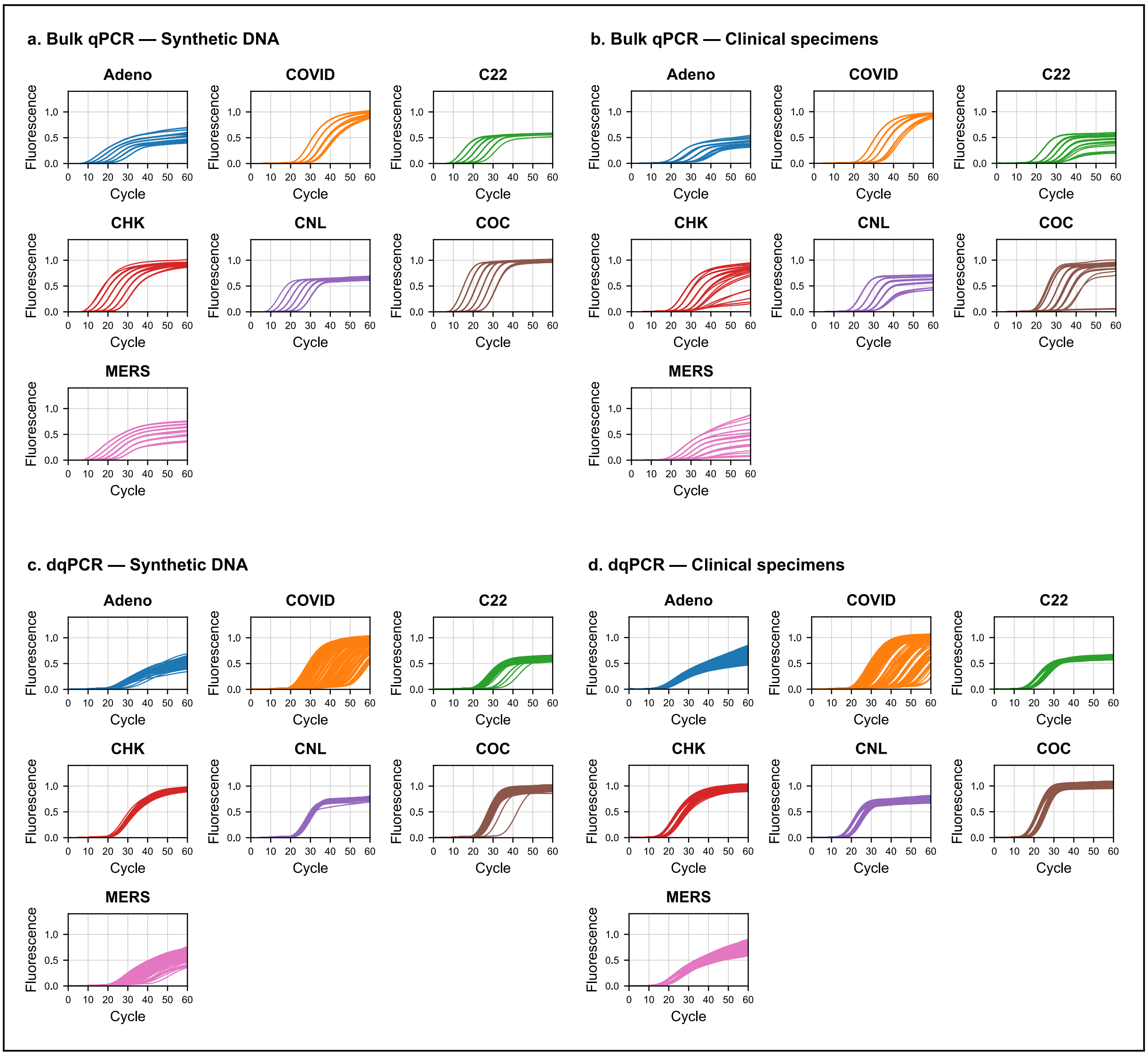


**Figure S3. AC distributions across experimental domains in the 7-plex respiratory virus panel.** ACs obtained for the 7-plex respiratory virus panel across four datasets: (a) bulk qPCR synthetic DNA (145 ACs obtained from one experiment are plotted, total dataset $n = 402$ ACs), (b) bulk qPCR contrived clinical specimens (153 ACs from one experiment, total dataset $n = 446$ ACs), (c) dqPCR synthetic DNA (11,132 ACs randomly selected from a total dataset with $n = 33,345$ ACs), and (d) dqPCR contrived clinical specimens (30,344 ACs randomly selected from a total dataset with $n = 59,237$ ACs). Bulk qPCR experiments were performed using decreasing concentrations of input DNA from $10^7$ copies to 1 copy + NTCs in triplicate. dqPCR experiments were performed at input concentrations of $10^6$, $10^5$, and $10^4$ copies + NTCs in triplicate.

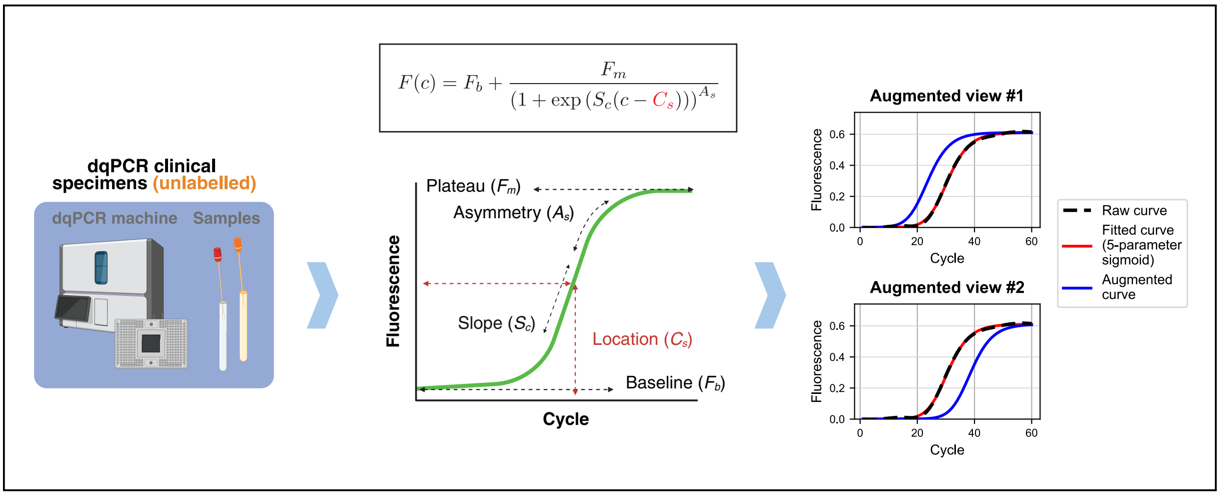


**Figure S4. Custom augmentation procedure for canonical ACs in dqPCR to capture concentration shifts of input DNA.**
Each raw AC from the dqPCR clinical dataset is fitted with a 5-parameter sigmoid model, and augmented views are generated by varying the location parameter ($C_s$) while keeping other parameters fixed. This procedure simulates realistic shifts in amplification timing associated with differences in input DNA concentration, while preserving the intrinsic target-specific kinetic and thermodynamic characteristics of the curve. Two independently augmented views of the same curve are used as positive pairs for self-supervised learning.

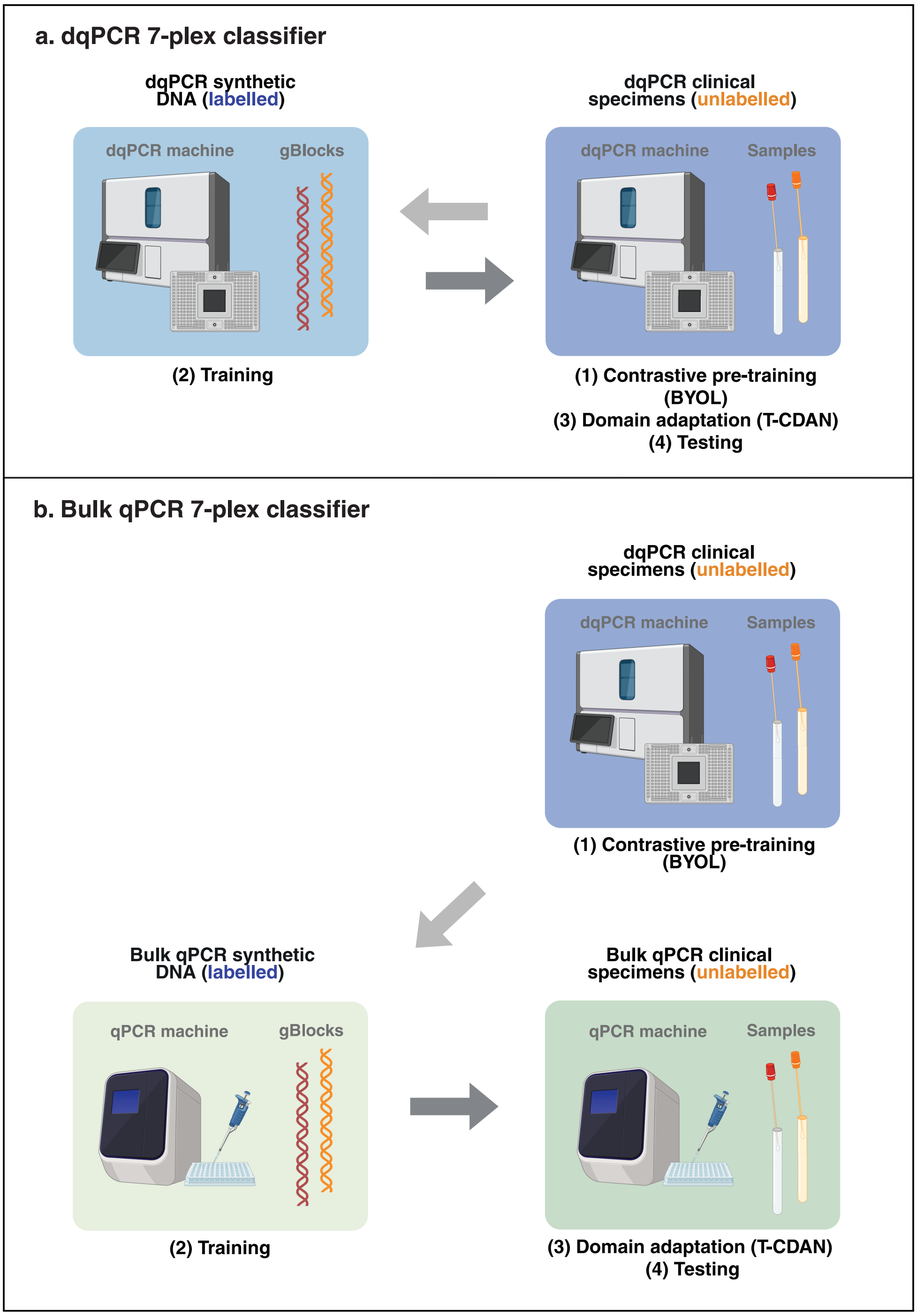


**Figure S5. Training and evaluation strategies for domain-adaptive classification of the 7-plex respiratory virus panel in digital and bulk qPCR.**

**a. dqPCR 7-plex classifier.** In the final classifier for the 7-plex single-channel respiratory viruses panel in dqPCR, the dqPCR contrived clinical specimens dataset (unlabelled) is used for self-supervised pretraining. The resulting representations are then used to train a classifier on the dqPCR synthetic DNA dataset (labelled), followed by domain adaptation. Classification performance is evaluated on the dqPCR contrived clinical specimens dataset. Figure created with BioRender®.

**b. Bulk qPCR 7-plex classifier.** In the final classifier for the 7-plex single-channel bulk qPCR assay, the dqPCR contrived clinical specimens dataset (unlabelled) is used for self-supervised pretraining. The model is then trained on the bulk qPCR synthetic DNA dataset (labelled) and further refined through unsupervised domain adaptation using the bulk qPCR contrived clinical specimens dataset as the target domain. Classification performance is evaluated on the bulk qPCR contrived clinical specimens dataset. Figure created with BioRender®.

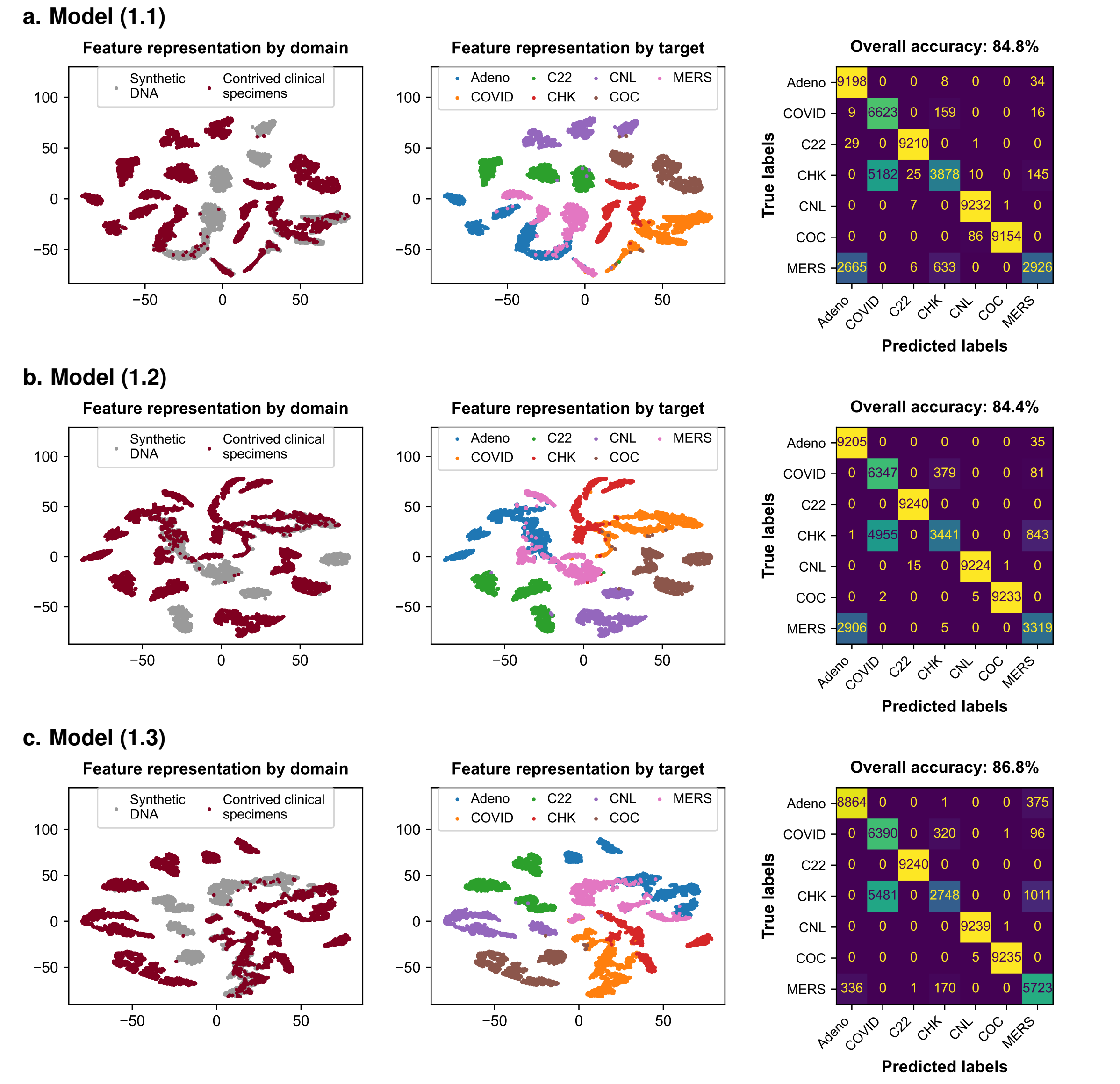


**d. Summary of models' architectures and classification performance**

| Model | Contrastive pretraining (BYOL) | Training | Domain Adaptation (T-CDAN) | Test | Accuracy (%) |
|---|---|---|---|---|---|
| Model (1.1) | – | dqPCR synthetic DNA | – | dqPCR contrived clinical specimens | 84.8 |
| Model (1.2) | – | dqPCR synthetic DNA | dqPCR contrived clinical specimens | dqPCR contrived clinical specimens | 84.4 |
| Model (1.3) | dqPCR contrived clinical specimens | dqPCR synthetic DNA | dqPCR contrived clinical specimens | dqPCR contrived clinical specimens | **86.8** |

**Figure S6. Self-supervised pretraining improves classification of dqPCR contrived clinical specimens.** Three models were evaluated: (1.1) a 1D-transformer trained on dqPCR synthetic DNA data, (1.2) T-CDAN for domain adaptation between synthetic DNA (source) and contrived clinical specimens (target) datasets, and (1.3) T-CDAN with prior self-supervised pretraining on dqPCR contrived clinical specimens using BYOL. For each model, feature representations are visualised by domain and by target class, alongside corresponding confusion matrices. Domain adaptation alone provides limited improvement, while self-supervised pretraining enhances feature alignment and class separability, leading to improved classification performance, with overall accuracy increasing from 84.8% to 86.8%.

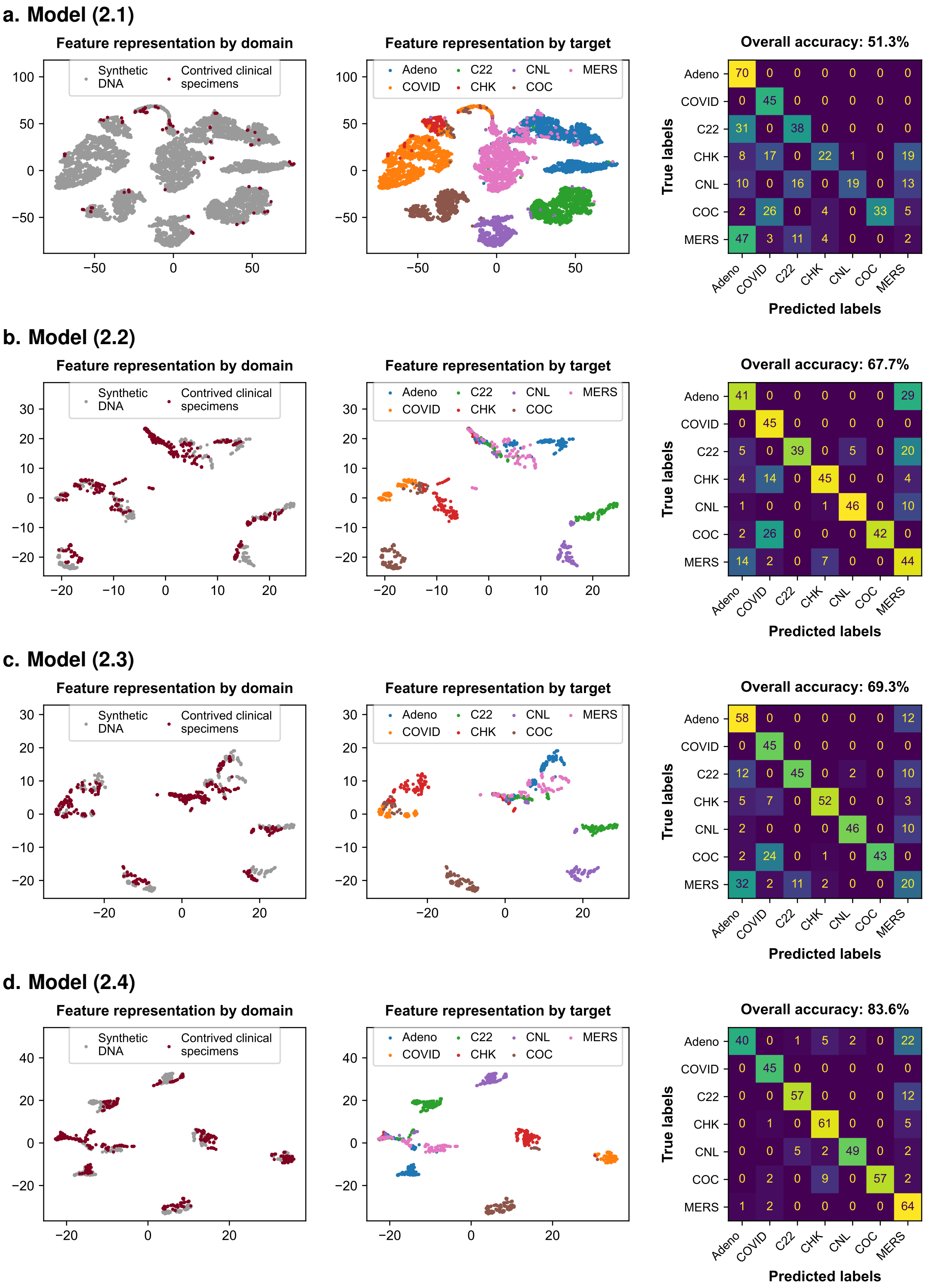


**e. Summary of models' architectures and classification performance**

| Model | Contrastive pretraining (BYOL) | Training | Domain Adaptation (T-CDAN) | Test set | Accuracy (%) |
|---|---|---|---|---|---|
| Model (2.1) | | dqPCR synthetic DNA | | Bulk qPCR contrived clinical specimens | 51.3 |
| Model (2.2) | | Bulk qPCR synthetic DNA | | Bulk qPCR contrived clinical specimens | 67.7 |
| Model (2.3) | | Bulk qPCR synthetic DNA | Bulk qPCR contrived clinical specimens | Bulk qPCR contrived clinical specimens | 69.3 |
| Model (2.4) | dqPCR contrived clinical specimens | Bulk qPCR synthetic DNA | Bulk qPCR contrived clinical specimens | Bulk qPCR contrived clinical specimens | **83.6** |

**Figure S7. Self-supervised pretraining and domain adaptation improve classification of bulk qPCR contrived clinical specimens.**

Four model configurations were evaluated: (2.1) a 1D-transformer trained on dqPCR synthetic DNA data and tested on bulk qPCR contrived clinical specimens data; (2.2) a 1D-transformer trained on bulk qPCR synthetic DNA data; (2.3) T-CDAN using the bulk qPCR synthetic DNA dataset as the labelled source domain and the bulk qPCR contrived clinical specimens dataset as the unlabelled target domain; and (2.4) T-CDAN with self-supervised pretraining (BYOL) on the dqPCR contrived clinical specimens dataset, followed by domain adaptation. Feature representations are visualised by domain and by target class, alongside confusion matrices. Classification accuracy improves from 51.3% (model 2.1) to 83.6% (model 2.4), demonstrating the combined benefit of domain adaptation and self-supervised pretraining for robust classification under domain shift.

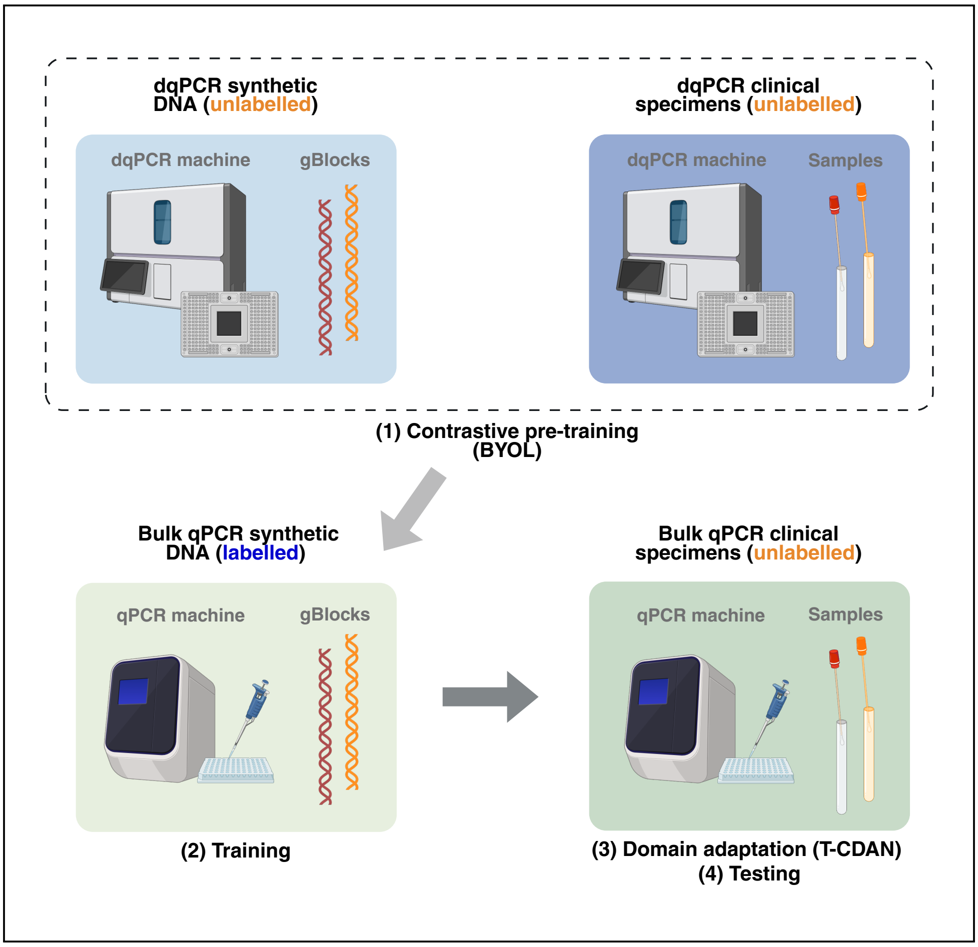


**Figure S8. Training and evaluation strategy for computational decoding of the stoichiometrically encoded 8-plex panel.**
In our final classifier for the 8-plex single-channel assay in bulk qPCR, the combined dqPCR synthetic DNA and contrived clinical specimens datasets (both unlabelled) were used for self-supervised pretraining. This serves as a basis for training on the bulk qPCR synthetic DNA dataset before domain adaptation using the bulk qPCR contrived clinical specimens dataset as the target domain. Classification accuracy is computed on the bulk qPCR contrived clinical specimens dataset. Figure created with BioRender®.
